## Supplementary Figure 1 for "An oligogenic architecture underlying ecological and reproductive divergence in sympatric populations"

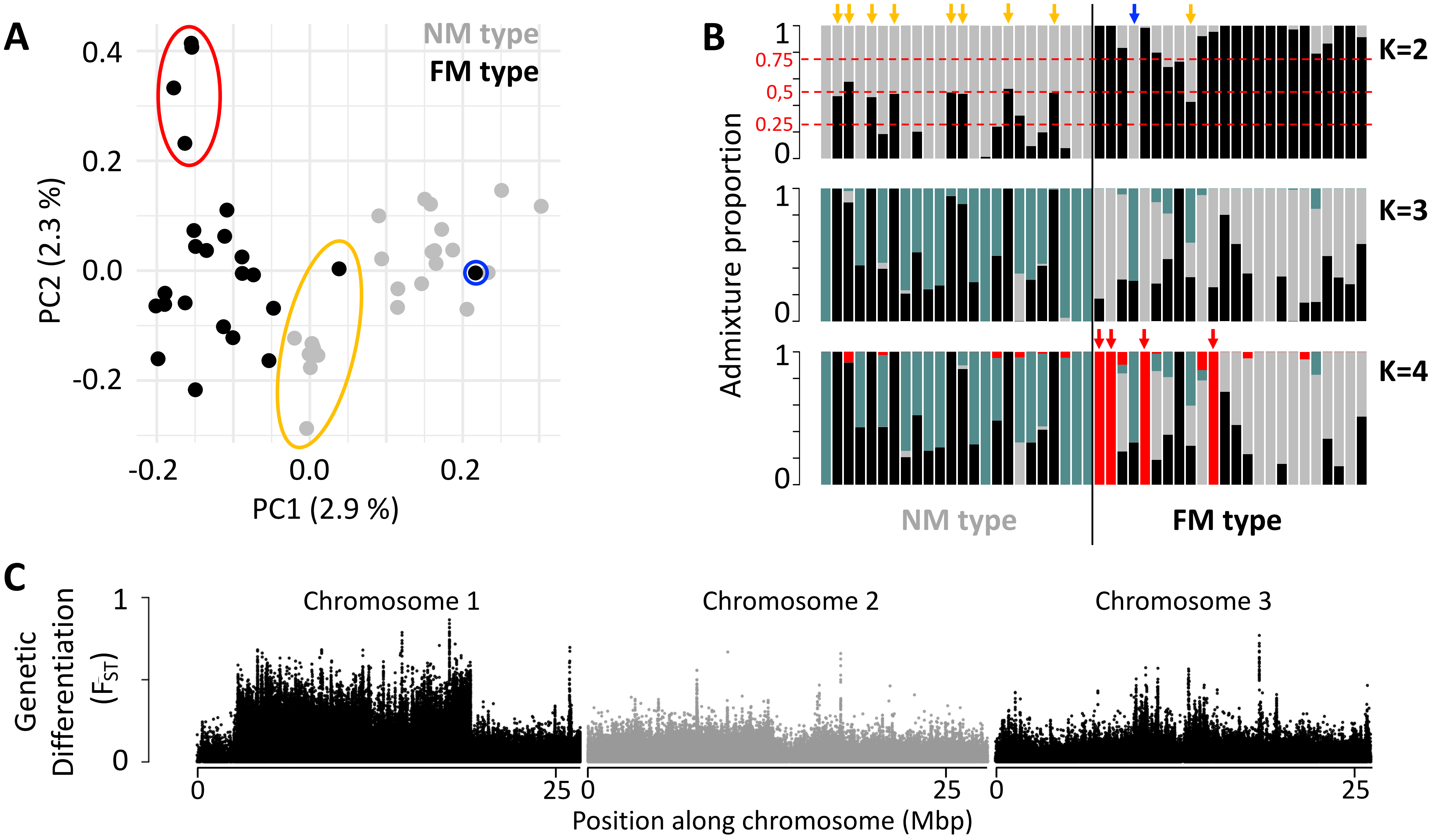


**Supplementary Figure 1 Population structure of the FM and NM types in Roscoff based on 721,000 genetic variants.**

(**A, B**) Principal component analysis (PCA; **A**) and admixture analysis (**B**) identify one migrant in time (blue; pure NM genotype caught at full moon), many potential F1 hybrids (yellow) and four individuals in the FM strain that appear genetically distict from all other samples (red). (**C**) Global genetic differentiation is limited, but there is a block of strong differentiation on chromosome 1.
