## Supplementary Figure 2 for "An oligogenic architecture underlying ecological and reproductive divergence in sympatric populations"

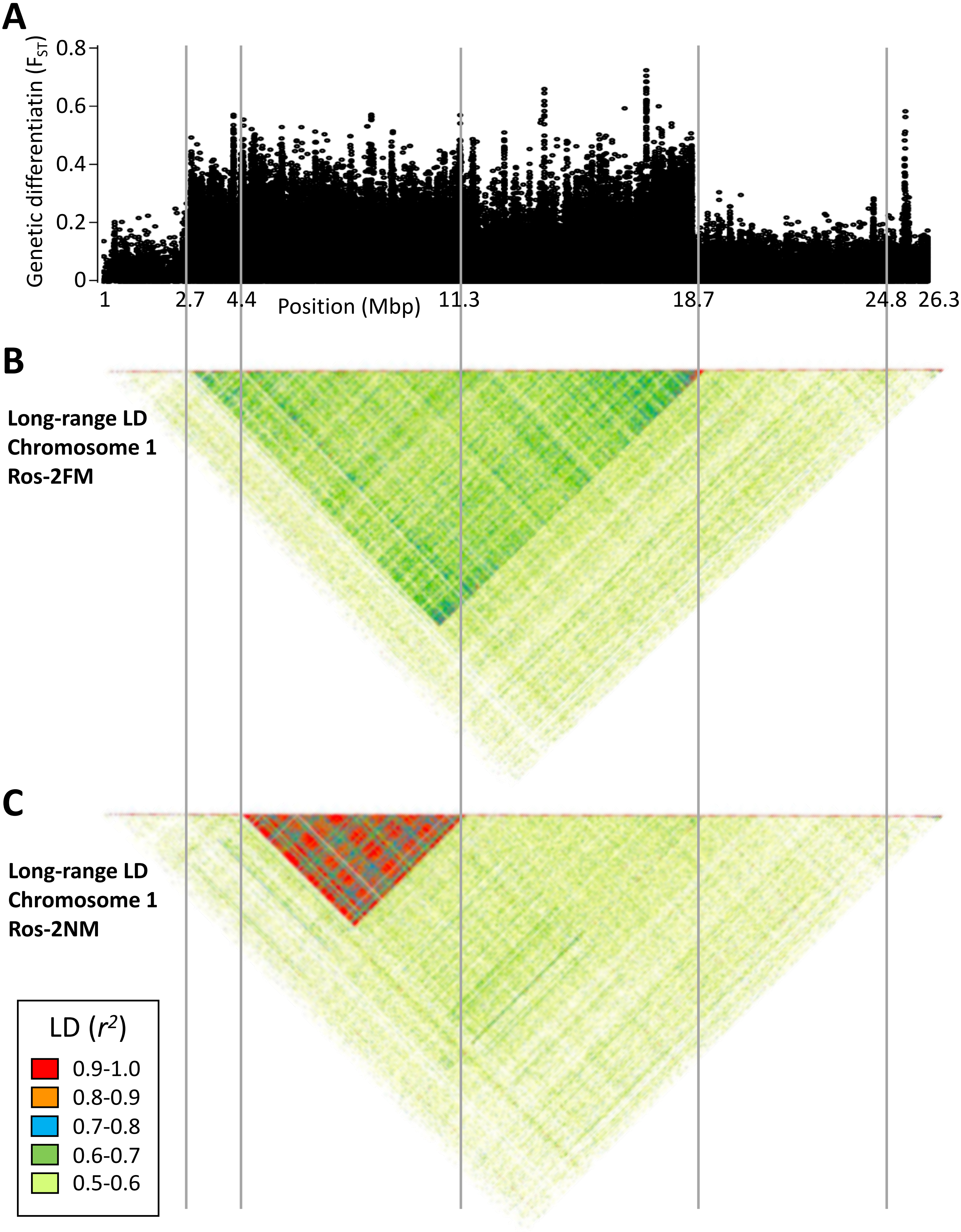


**Supplementary Figure 2 Long range linkage disequilibrium along chromosome 1**. Blocks of elevated genetic differentiation (**A**) correspond to blocks of elevated long-range LD in the FM type (**B**) and the NM type (**C**). Elevated long-range LD suggets that the inversion is polymorphic in the respective population.
