## Supplementary Figure 3 for "An oligogenic architecture underlying ecological and reproductive divergence in sympatric populations"

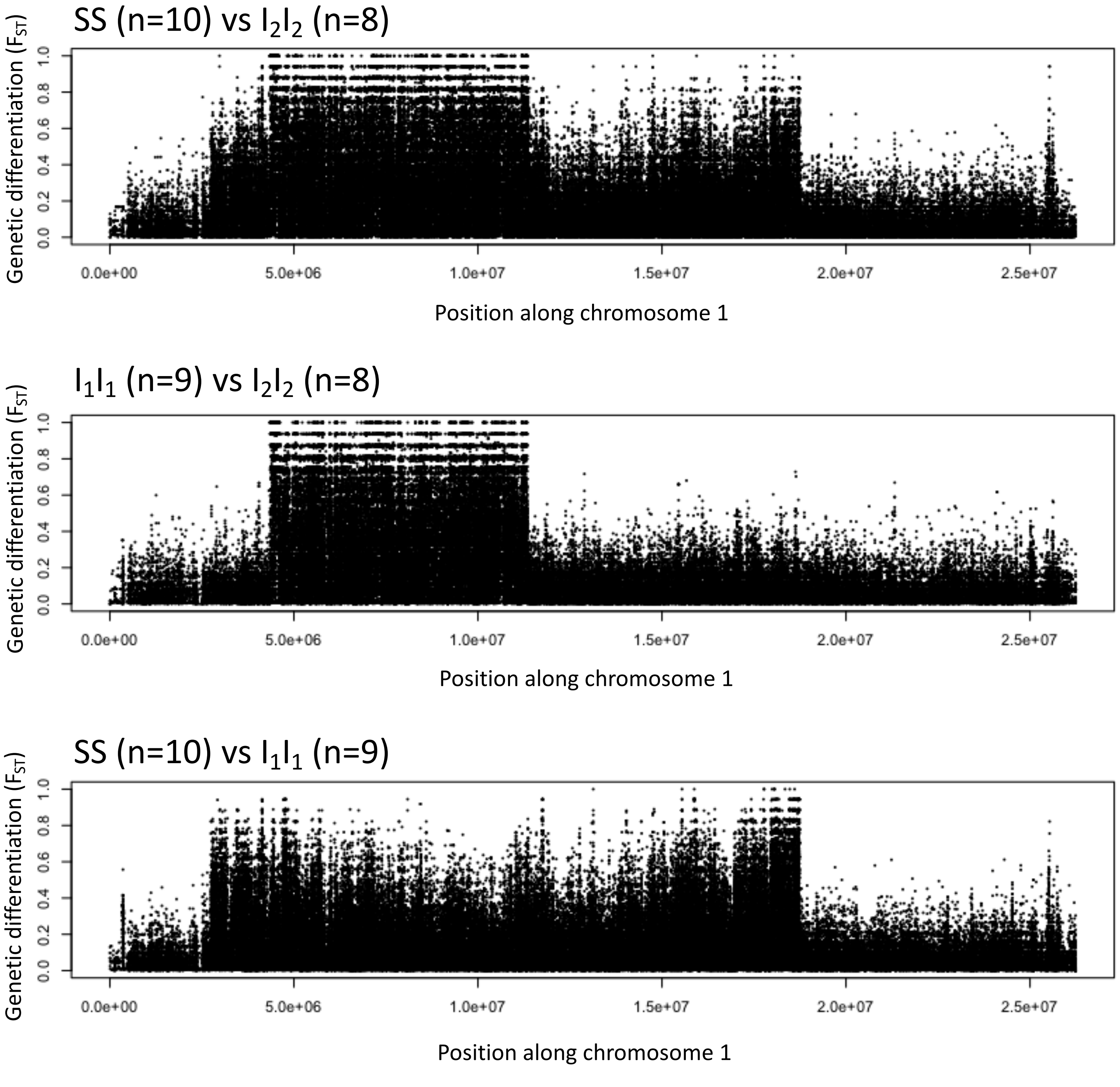


**Supplementary Figure 3** Genetic differentiation between homozygotes of the standard haplotype (SS) and inversion haplotypes ($I_1I_1$, $I_2I_2$).
