## Supplementary Figure 4 for "An oligogenic architecture underlying ecological and reproductive divergence in sympatric populations"

### Supplementary Figure 4 Windowed PCA along chromosome 1 (500kb sliding windows, 100kb steps).

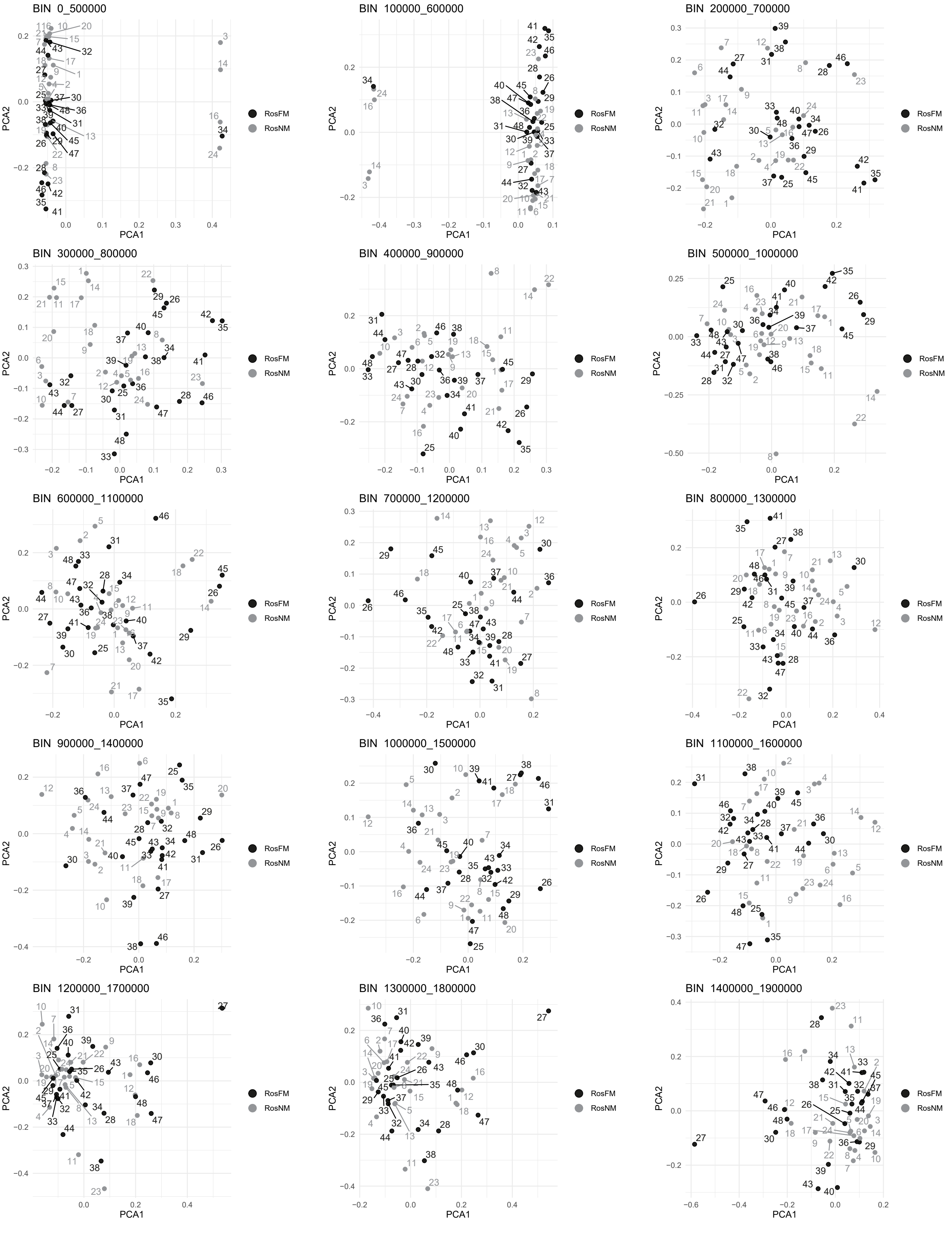

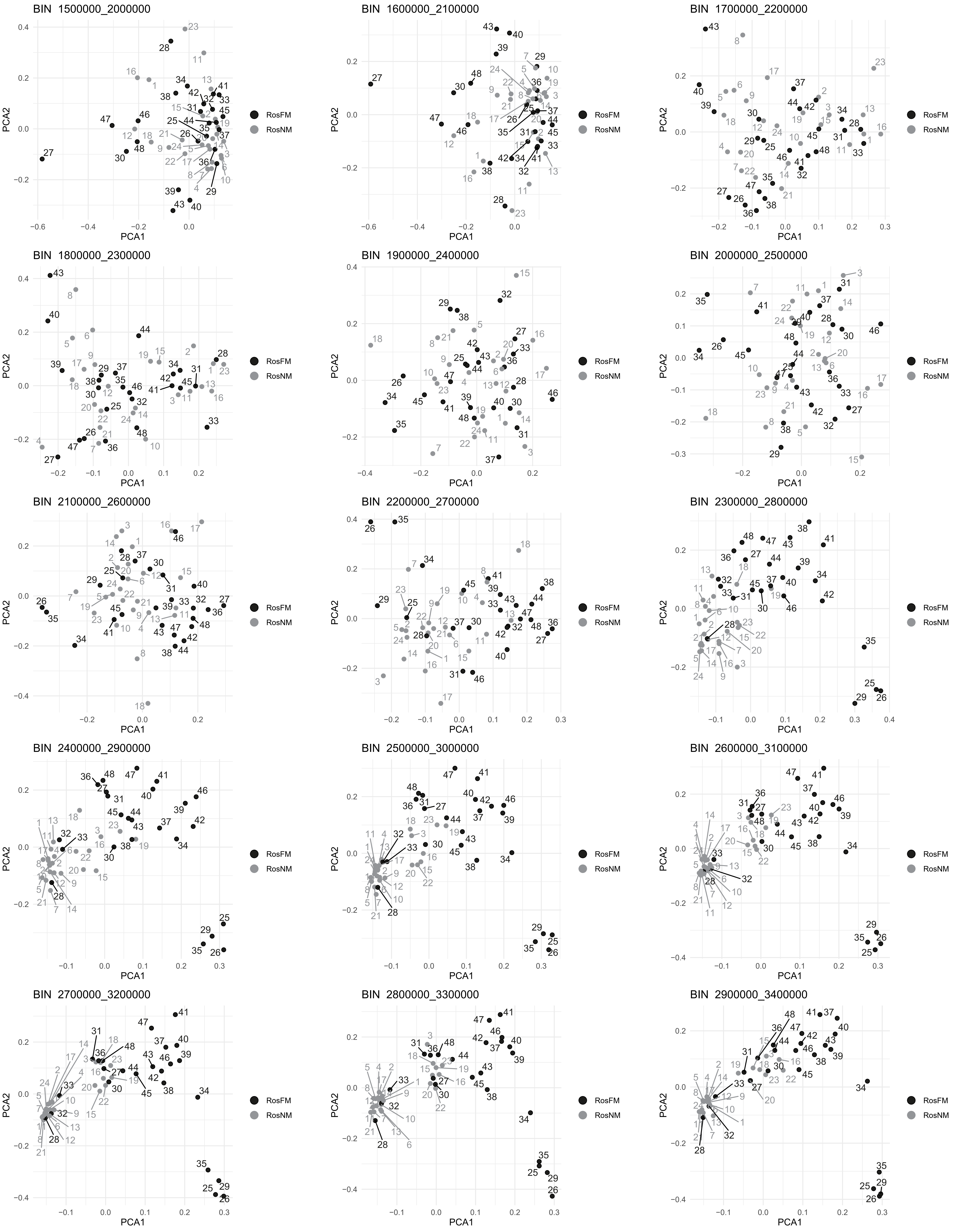
BIN 1500000_2000000
BIN 1600000_2100000
BIN 1700000_2200000
BIN 1800000_2300000
BIN 1900000_2400000
BIN 2000000_2500000
BIN 2100000_2600000
BIN 2200000_2700000
BIN 2300000_2800000
BIN 2400000_2900000
BIN 2500000_3000000
BIN 2600000_3100000
BIN 2700000_3200000
BIN 2800000_3300000
BIN 2900000_3400000
PCA1
PCA2
RosFM
RosNM

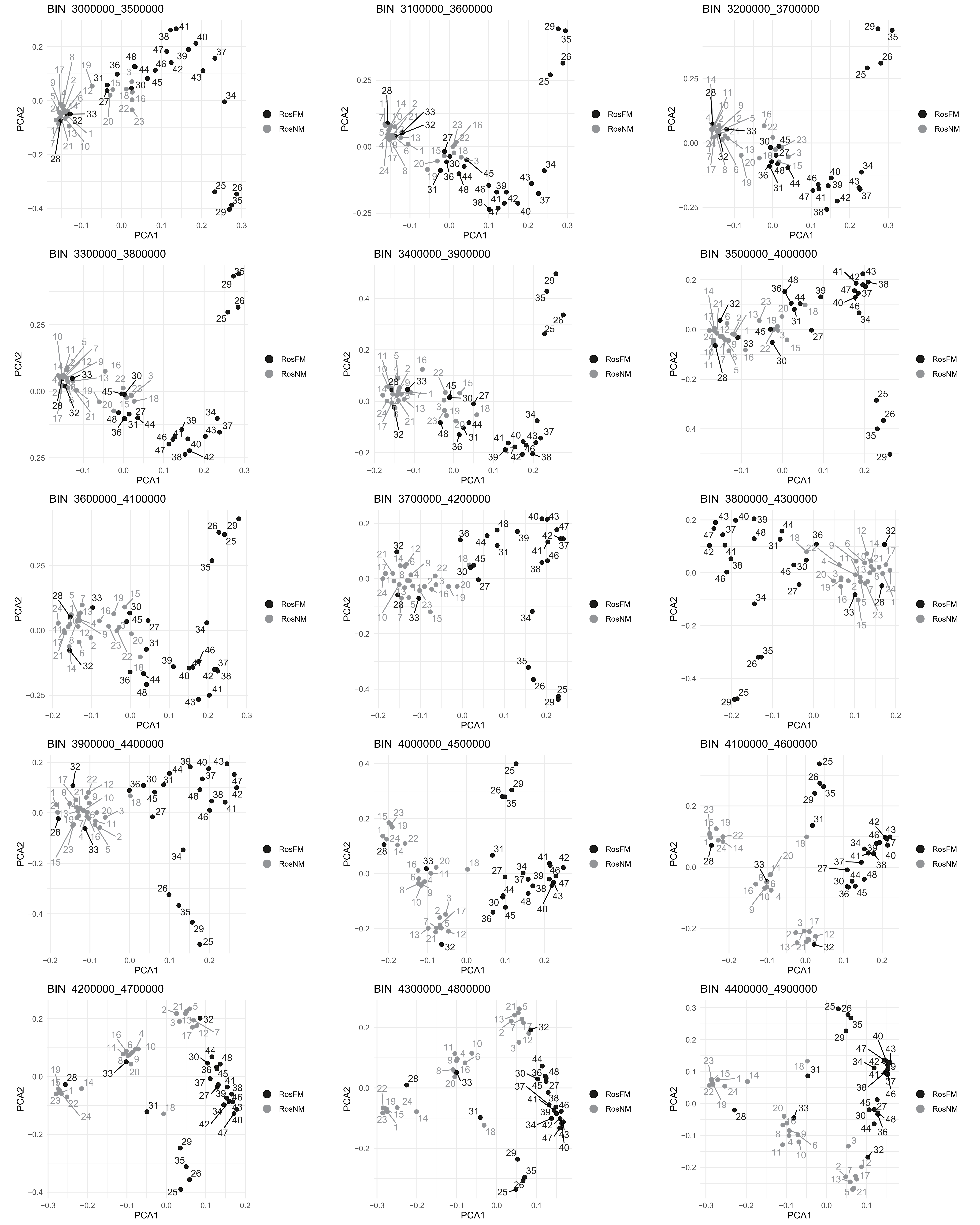
BIN 3000000_3500000
BIN 3100000_3600000
BIN 3200000_3700000
BIN 3300000_3800000
BIN 3400000_3900000
BIN 3500000_4000000
BIN 3600000_4100000
BIN 3700000_4200000
BIN 3800000_4300000
BIN 3900000_4400000
BIN 4000000_4500000
BIN 4100000_4600000
BIN 4200000_4700000
BIN 4300000_4800000
BIN 4400000_4900000
PCA1
PCA2
RosFM
RosNM

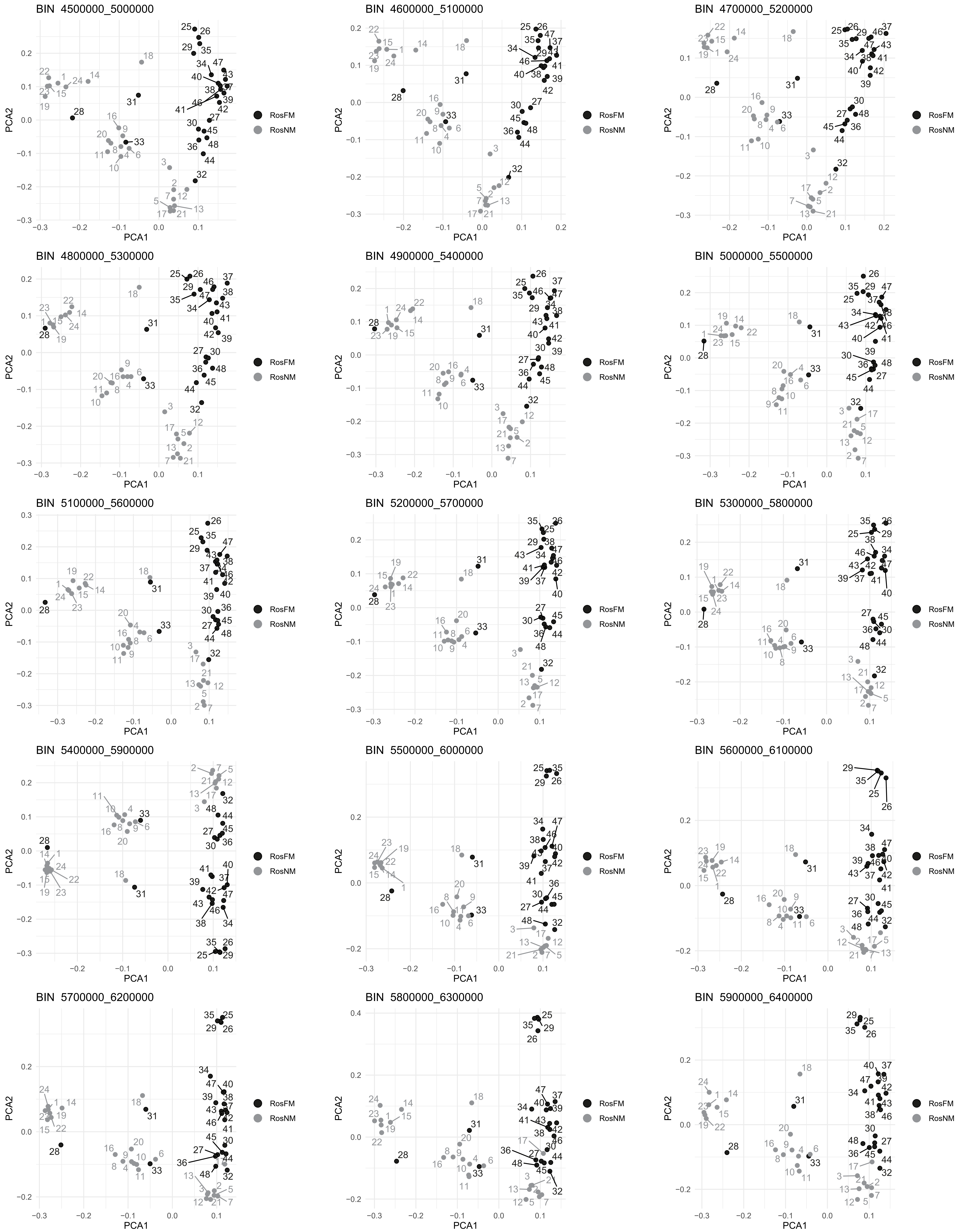
BIN 4500000_5000000
BIN 4600000_5100000
BIN 4700000_5200000
BIN 4800000_5300000
BIN 4900000_5400000
BIN 5000000_5500000
BIN 5100000_5600000
BIN 5200000_5700000
BIN 5300000_5800000
BIN 5400000_5900000
BIN 5500000_6000000
BIN 5600000_6100000
BIN 5700000_6200000
BIN 5800000_6300000
BIN 5900000_6400000
PCA1
PCA2
RosFM
RosNM

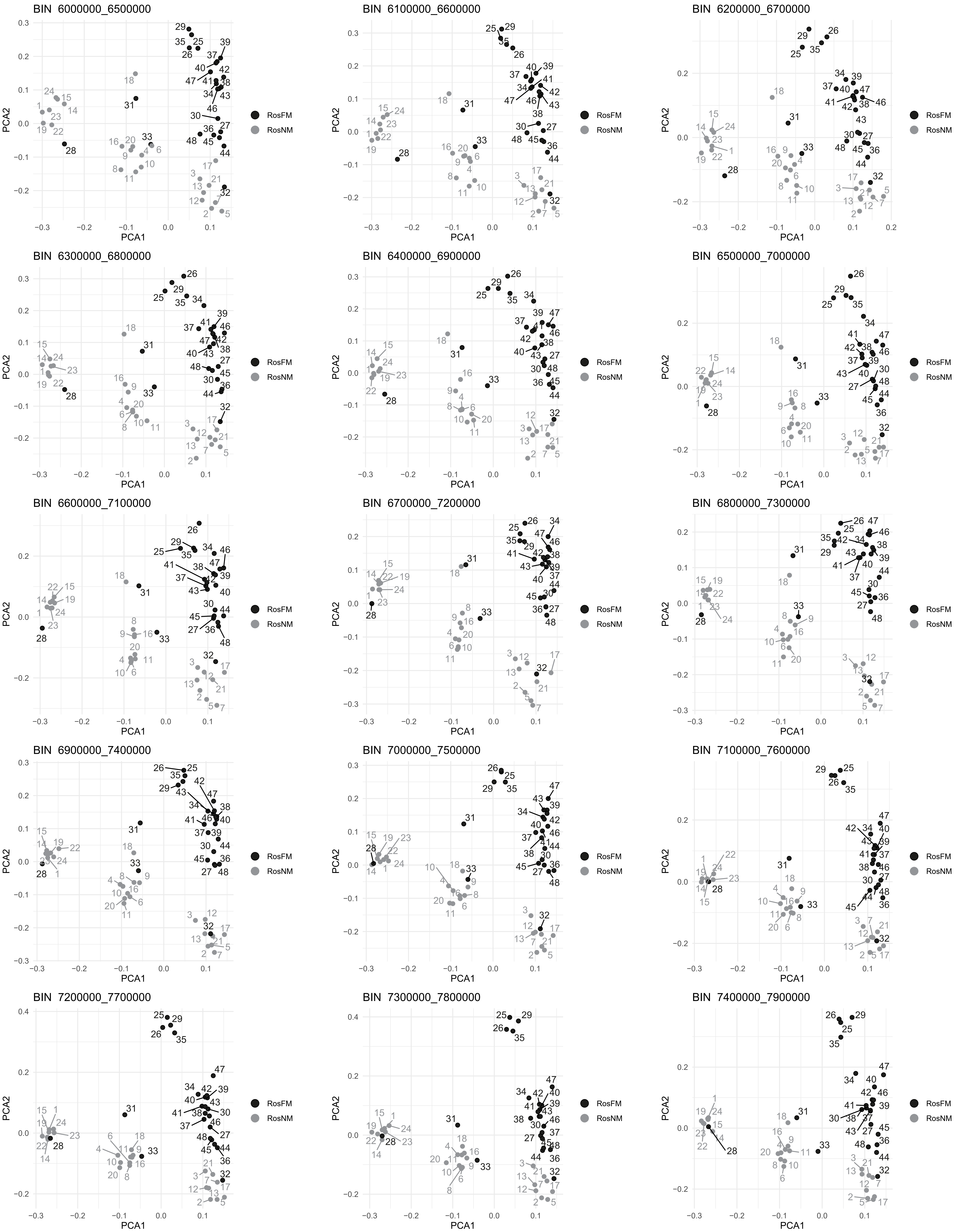
BIN 6000000_6500000
BIN 6100000_6600000
BIN 6200000_6700000
BIN 6300000_6800000
BIN 6400000_6900000
BIN 6500000_7000000
BIN 6600000_7100000
BIN 6700000_7200000
BIN 6800000_7300000
BIN 6900000_7400000
BIN 7000000_7500000
BIN 7100000_7600000
BIN 7200000_7700000
BIN 7300000_7800000
BIN 7400000_7900000
PCA1
PCA2
RosFM
RosNM

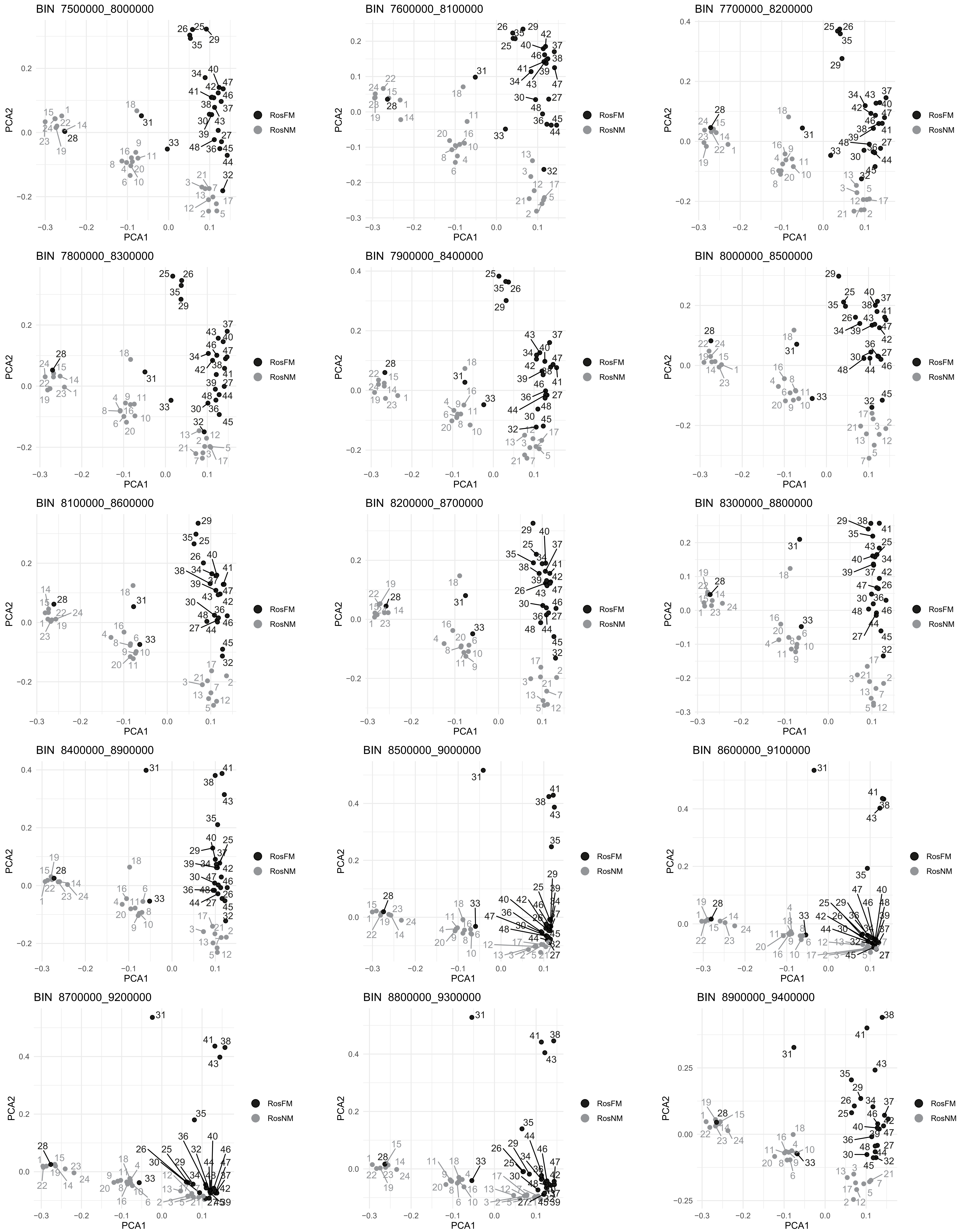
BIN 7500000_8000000
BIN 7600000_8100000
BIN 7700000_8200000
BIN 7800000_8300000
BIN 7900000_8400000
BIN 8000000_8500000
BIN 8100000_8600000
BIN 8200000_8700000
BIN 8300000_8800000
BIN 8400000_8900000
BIN 8500000_9000000
BIN 8600000_9100000
BIN 8700000_9200000
BIN 8800000_9300000
BIN 8900000_9400000
PCA1
PCA2
RosFM
RosNM

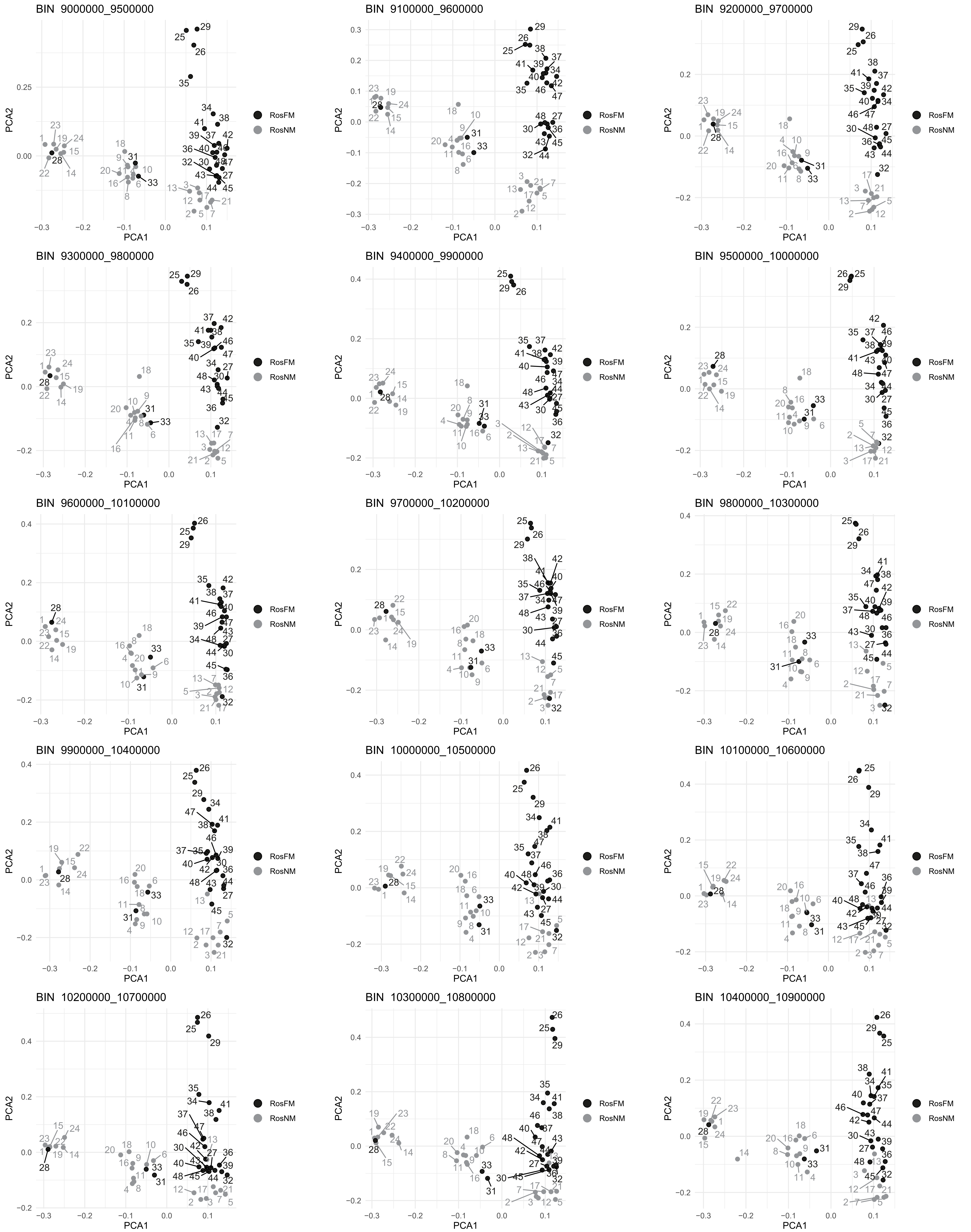

BIN 9000000_9500000
BIN 9100000_9600000
BIN 9200000_9700000
BIN 9300000_9800000
BIN 9400000_9900000
BIN 9500000_10000000
BIN 9600000_10100000
BIN 9700000_10200000
BIN 9800000_10300000
BIN 9900000_10400000
BIN 10000000_10500000
BIN 10100000_10600000
BIN 10200000_10700000
BIN 10300000_10800000
BIN 10400000_10900000
PCA1
PCA2
RosFM
RosNM

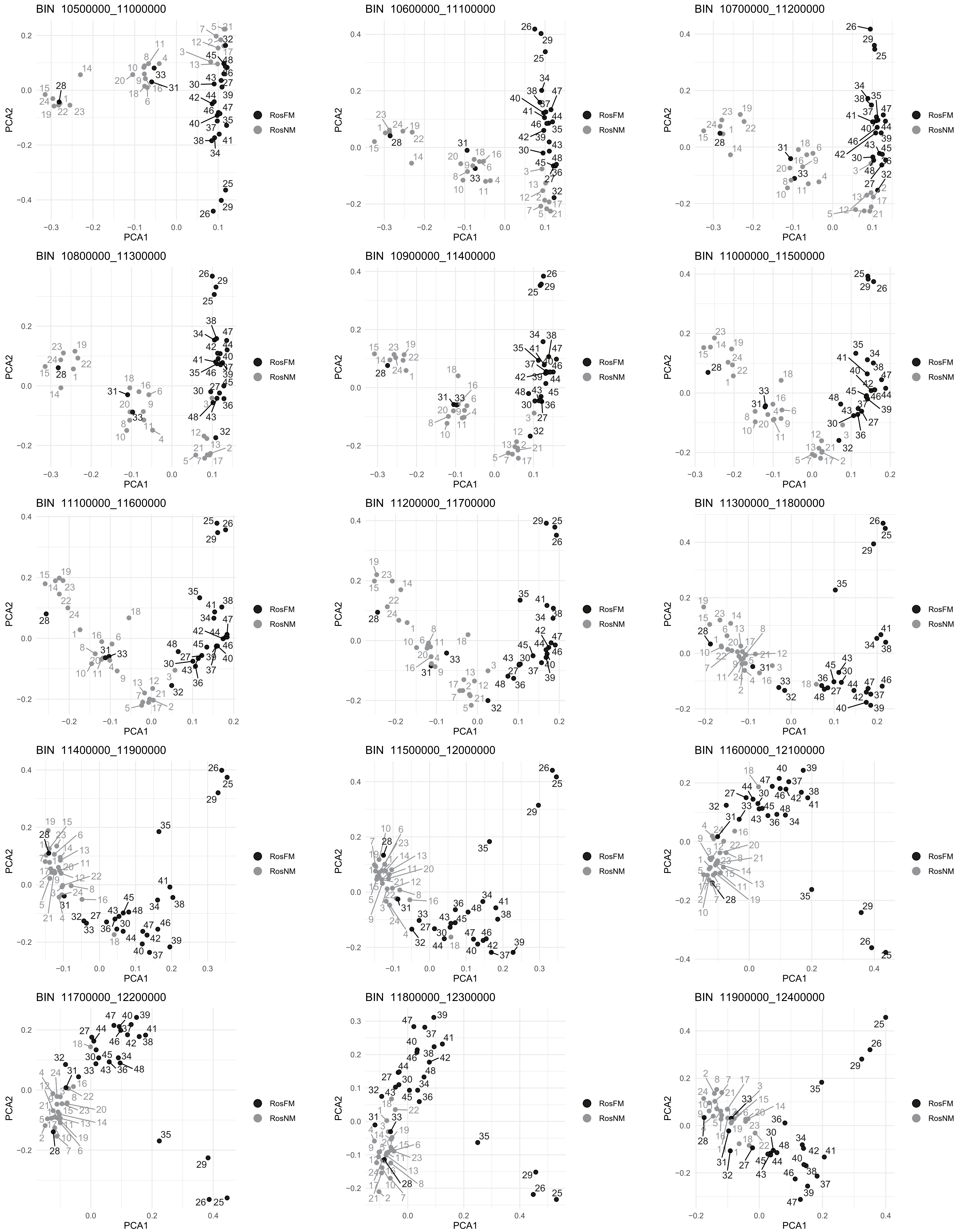
BIN 10500000_11000000
BIN 10600000_11100000
BIN 10700000_11200000
BIN 10800000_11300000
BIN 10900000_11400000
BIN 11000000_11500000
BIN 11100000_11600000
BIN 11200000_11700000
BIN 11300000_11800000
BIN 11400000_11900000
BIN 11500000_12000000
BIN 11600000_12100000
BIN 11700000_12200000
BIN 11800000_12300000
BIN 11900000_12400000
PCA1
PCA2
RosFM
RosNM

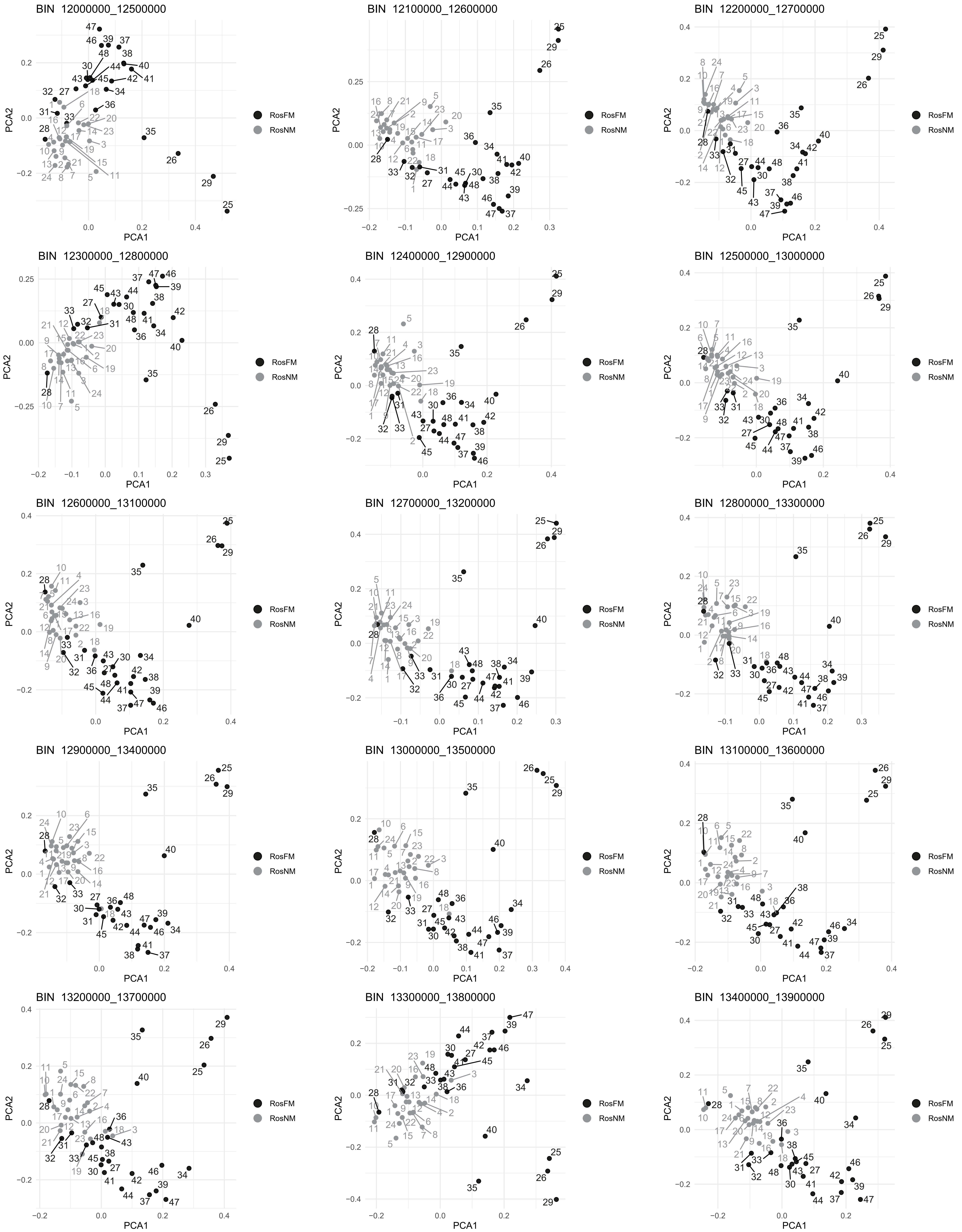
BIN 12000000_12500000
BIN 12100000_12600000
BIN 12200000_12700000
BIN 12300000_12800000
BIN 12400000_12900000
BIN 12500000_13000000
BIN 12600000_13100000
BIN 12700000_13200000
BIN 12800000_13300000
BIN 12900000_13400000
BIN 13000000_13500000
BIN 13100000_13600000
BIN 13200000_13700000
BIN 13300000_13800000
BIN 13400000_13900000
PCA1
PCA2
RosFM
RosNM

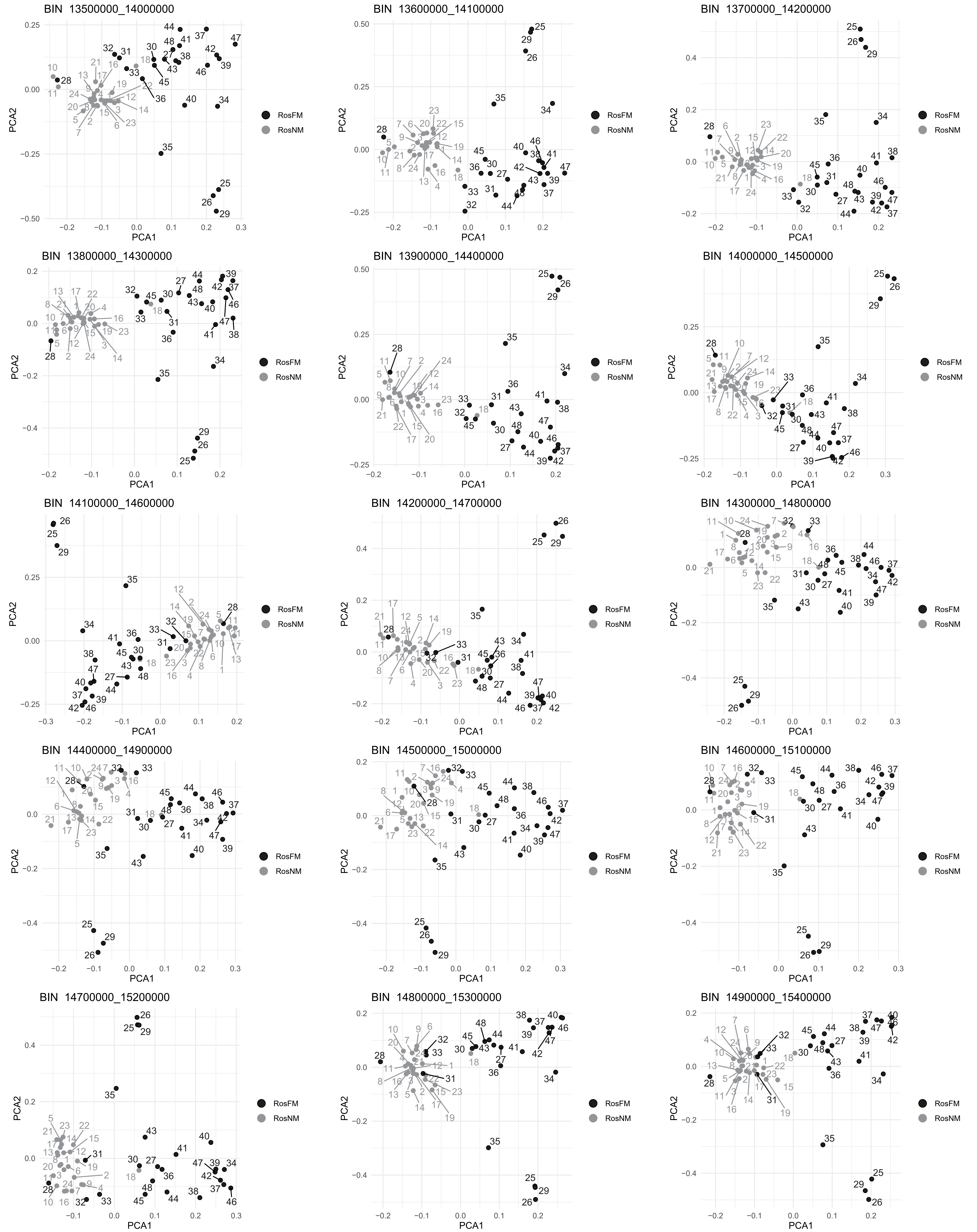
BIN 13500000_14000000
BIN 13600000_14100000
BIN 13700000_14200000
BIN 13800000_14300000
BIN 13900000_14400000
BIN 14000000_14500000
BIN 14100000_14600000
BIN 14200000_14700000
BIN 14300000_14800000
BIN 14400000_14900000
BIN 14500000_15000000
BIN 14600000_15100000
BIN 14700000_15200000
BIN 14800000_15300000
BIN 14900000_15400000
PCA1
PCA2
RosFM
RosNM

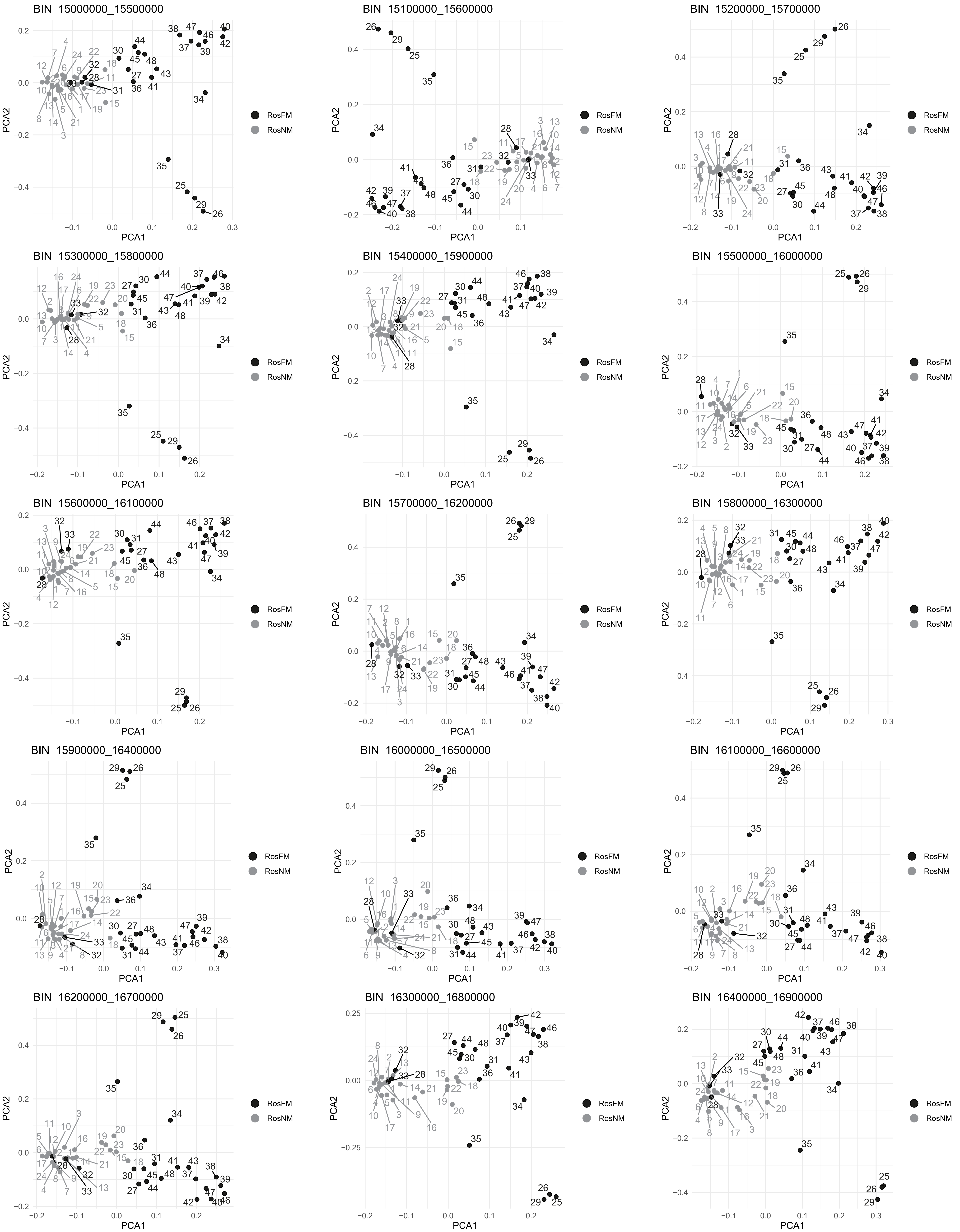
BIN 15000000_15500000
BIN 15100000_15600000
BIN 15200000_15700000
BIN 15300000_15800000
BIN 15400000_15900000
BIN 15500000_16000000
BIN 15600000_16100000
BIN 15700000_16200000
BIN 15800000_16300000
BIN 15900000_16400000
BIN 16000000_16500000
BIN 16100000_16600000
BIN 16200000_16700000
BIN 16300000_16800000
BIN 16400000_16900000
PCA1
PCA2
RosFM
RosNM

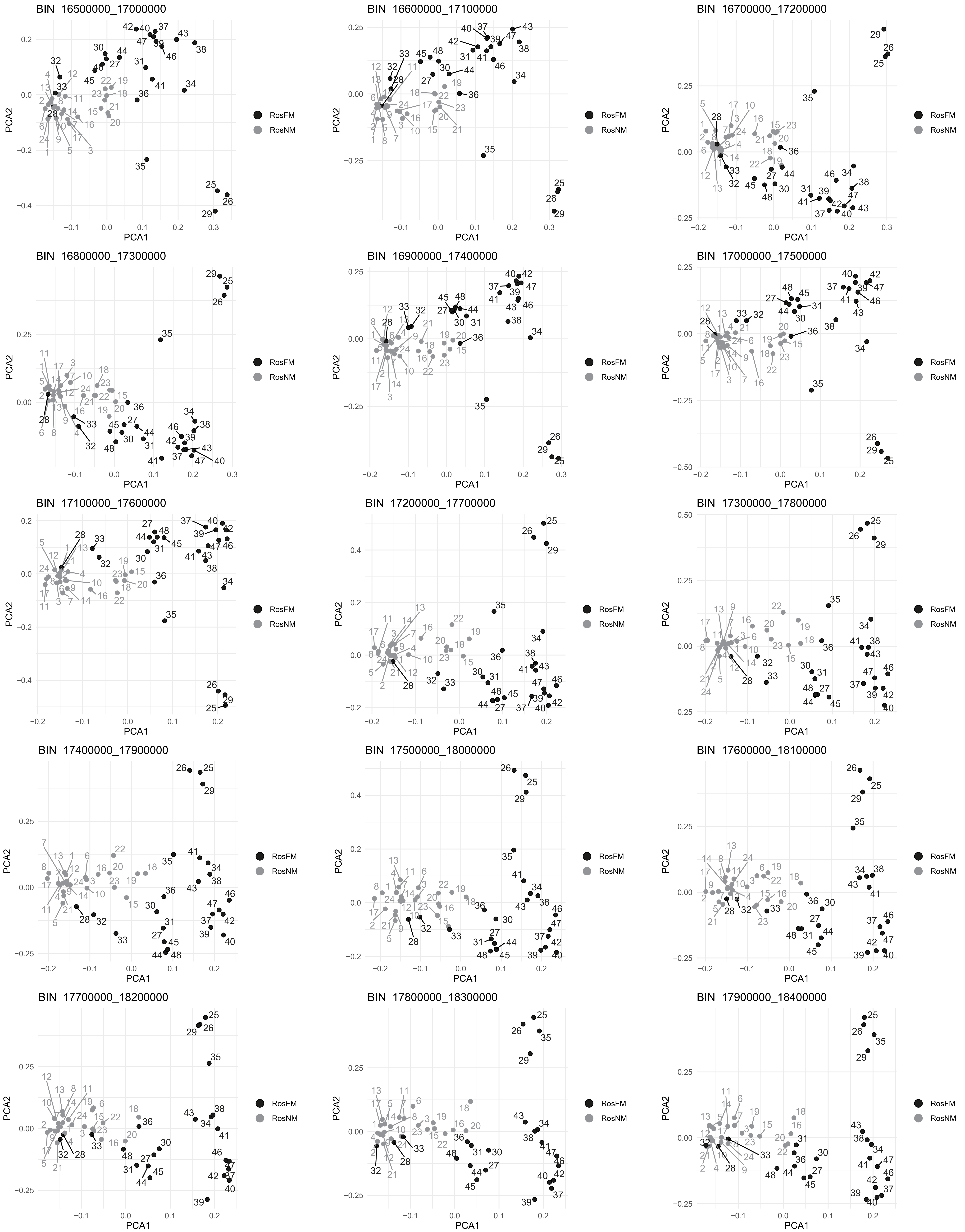
BIN 16500000_17000000
BIN 16600000_17100000
BIN 16700000_17200000
BIN 16800000_17300000
BIN 16900000_17400000
BIN 17000000_17500000
BIN 17100000_17600000
BIN 17200000_17700000
BIN 17300000_17800000
BIN 17400000_17900000
BIN 17500000_18000000
BIN 17600000_18100000
BIN 17700000_18200000
BIN 17800000_18300000
BIN 17900000_18400000
PCA1
PCA2
RosFM
RosNM

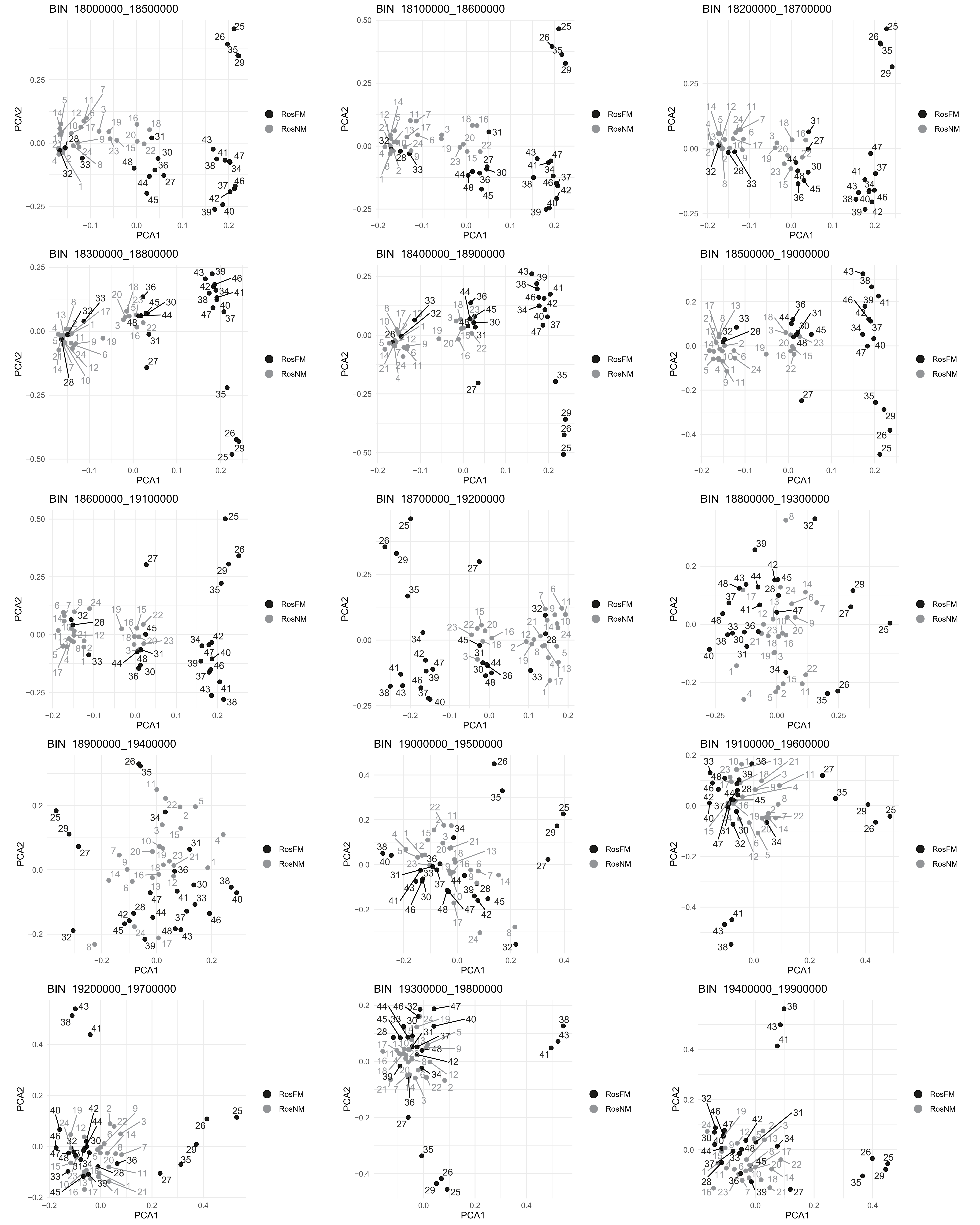
BIN 18000000_18500000
BIN 18100000_18600000
BIN 18200000_18700000
BIN 18300000_18800000
BIN 18400000_18900000
BIN 18500000_19000000
BIN 18600000_19100000
BIN 18700000_19200000
BIN 18800000_19300000
BIN 18900000_19400000
BIN 19000000_19500000
BIN 19100000_19600000
BIN 19200000_19700000
BIN 19300000_19800000
BIN 19400000_19900000
PCA1
PCA2
RosFM
RosNM

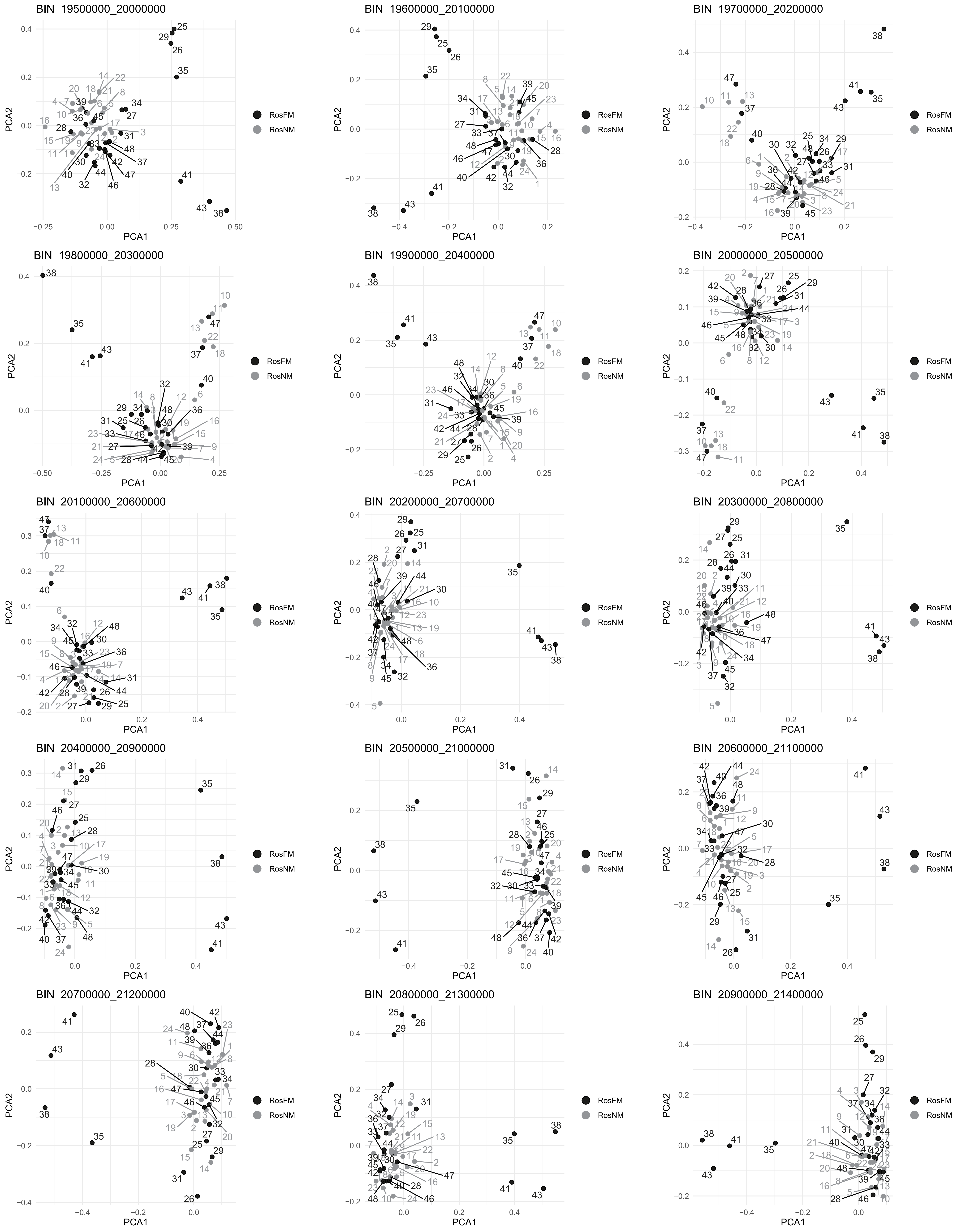
BIN 19500000_20000000
BIN 19600000_20100000
BIN 19700000_20200000
BIN 19800000_20300000
BIN 19900000_20400000
BIN 20000000_20500000
BIN 20100000_20600000
BIN 20200000_20700000
BIN 20300000_20800000
BIN 20400000_20900000
BIN 20500000_21000000
BIN 20600000_21100000
BIN 20700000_21200000
BIN 20800000_21300000
BIN 20900000_21400000
PCA1
PCA2
RosFM
RosNM

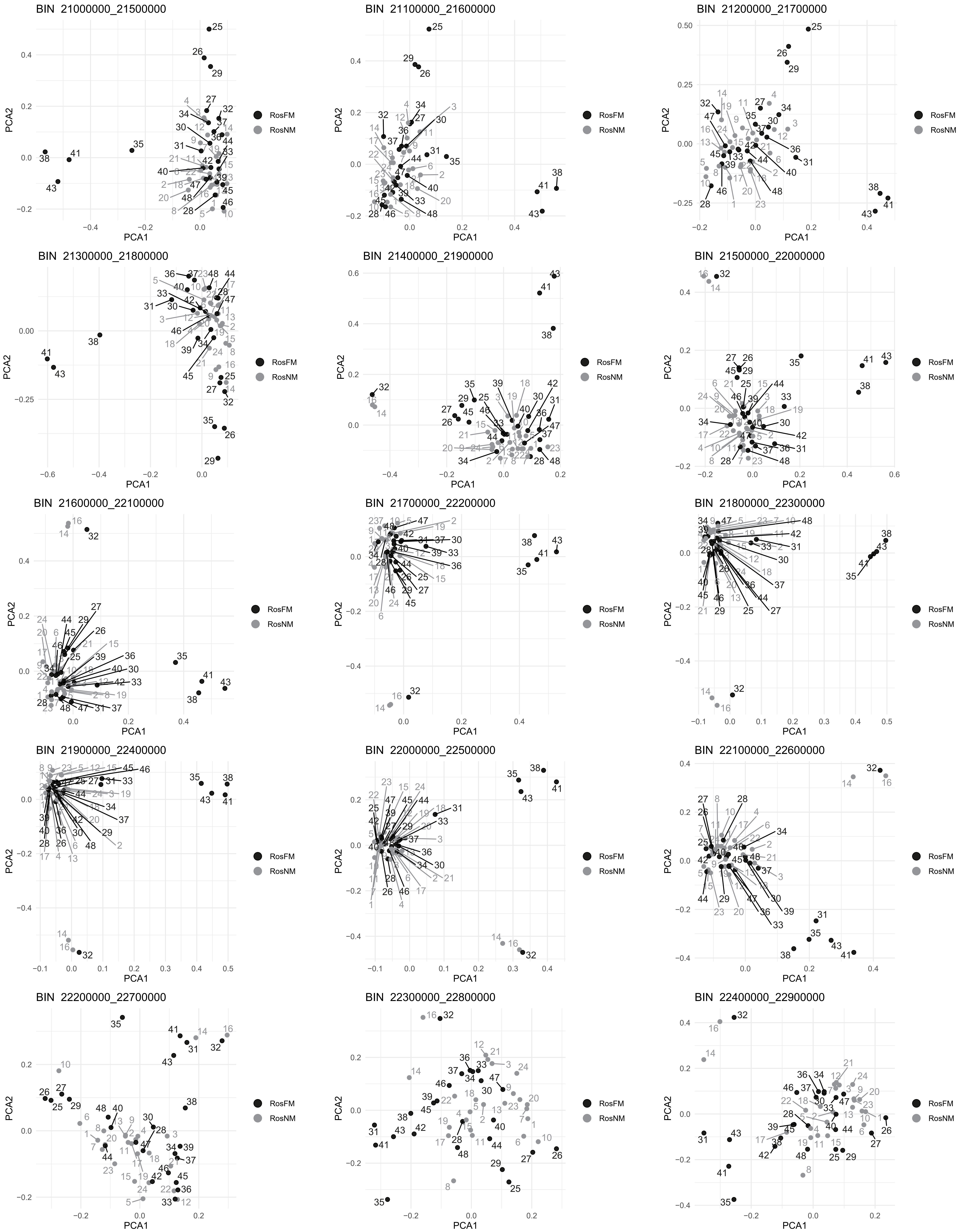
BIN 21000000_21500000
BIN 21100000_21600000
BIN 21200000_21700000
BIN 21300000_21800000
BIN 21400000_21900000
BIN 21500000_22000000
BIN 21600000_22100000
BIN 21700000_22200000
BIN 21800000_22300000
BIN 21900000_22400000
BIN 22000000_22500000
BIN 22100000_22600000
BIN 22200000_22700000
BIN 22300000_22800000
BIN 22400000_22900000
PCA1
PCA2
RosFM
RosNM

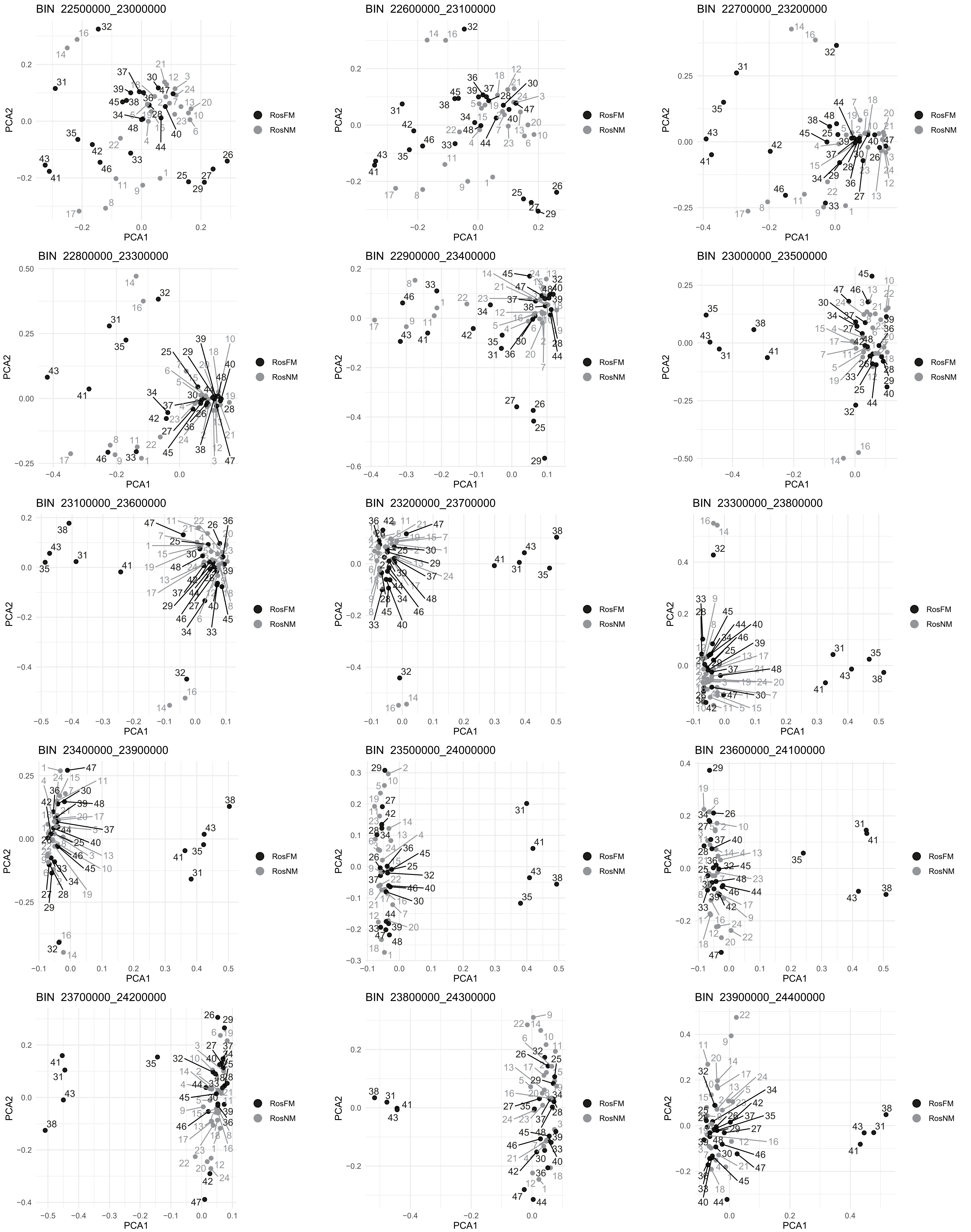
BIN 22500000_23000000
BIN 22600000_23100000
BIN 22700000_23200000
BIN 22800000_23300000
BIN 22900000_23400000
BIN 23000000_23500000
BIN 23100000_23600000
BIN 23200000_23700000
BIN 23300000_23800000
BIN 23400000_23900000
BIN 23500000_24000000
BIN 23600000_24100000
BIN 23700000_24200000
BIN 23800000_24300000
BIN 23900000_24400000
PCA1
PCA2
RosFM
RosNM

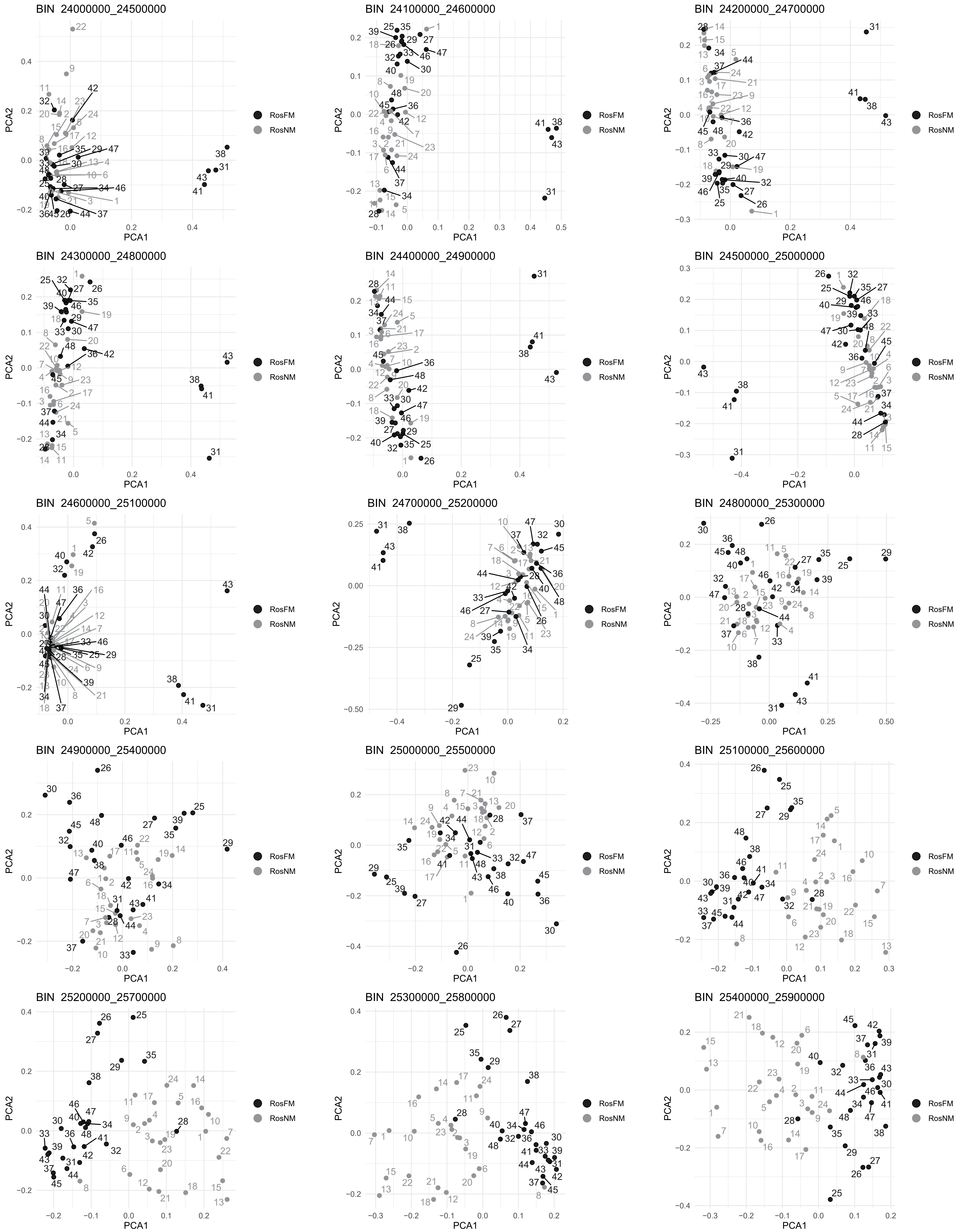
BIN 24000000_24500000
BIN 24100000_24600000
BIN 24200000_24700000
BIN 24300000_24800000
BIN 24400000_24900000
BIN 24500000_25000000
BIN 24600000_25100000
BIN 24700000_25200000
BIN 24800000_25300000
BIN 24900000_25400000
BIN 25000000_25500000
BIN 25100000_25600000
BIN 25200000_25700000
BIN 25300000_25800000
BIN 25400000_25900000
PCA1
PCA2
RosFM
RosNM

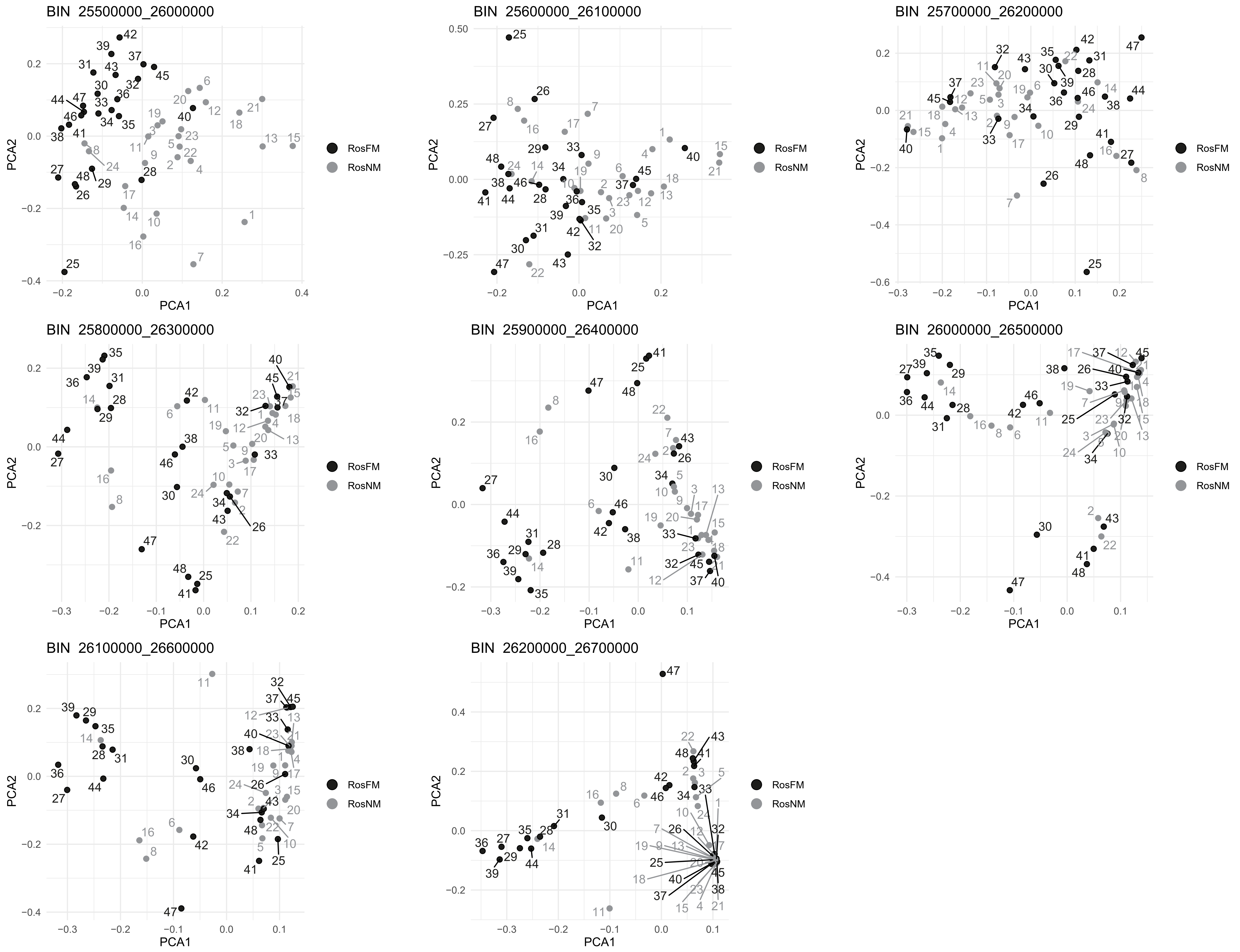
BIN 25500000_26000000
BIN 25600000_26100000
BIN 25700000_26200000
BIN 25800000_26300000
BIN 25900000_26400000
BIN 26000000_26500000
BIN 26100000_26600000
BIN 26200000_26700000
PCA1
PCA2
RosFM
RosNM
