## Supplementary Figure 5 for "An oligogenic architecture underlying ecological and reproductive divergence in sympatric populations"

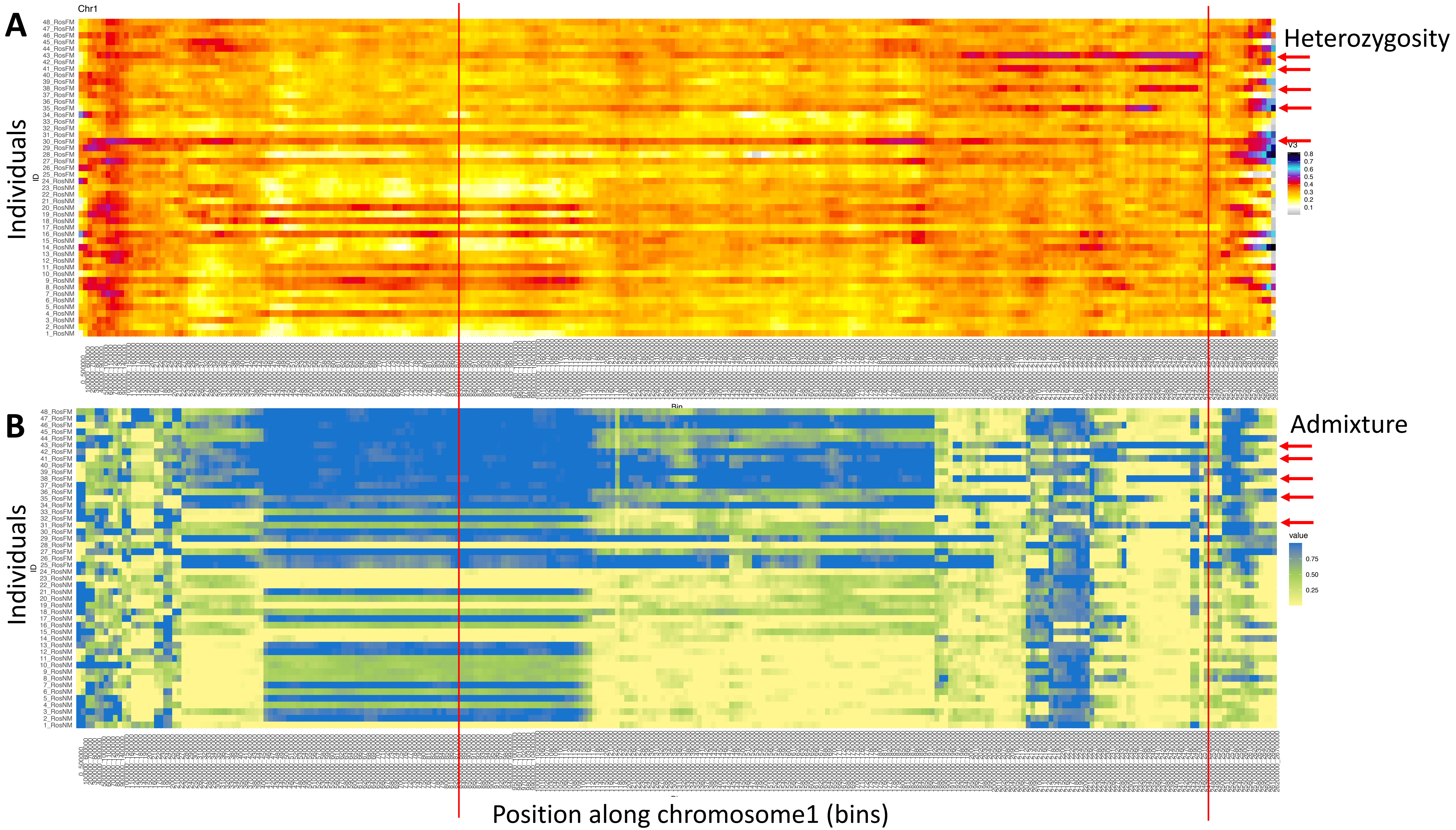


**Supplementary Figure 5 Windowed analysis for observed heterozygosity (A) and Admixture (B) along chromosome 1.**
(**A**) A number of individuals (red arrows) show elevated heterozygosity beyond the end of In(1a). In three of them (individuals 38, 41 and 43) the ends of that region coincide (red lines). These individuals are considered $SI_3$ heterozygotes, carrying In(1c). In the other two individuals the region is shorter or longer, possibly pointing to yet other inversions. (**B**) Admixture scores from windowed analysis support the same individuals to be genetically distinct in the right arm of In(1c).
