## Supplementary Figure 6 for "An oligogenic architecture underlying ecological and reproductive divergence in sympatric populations"

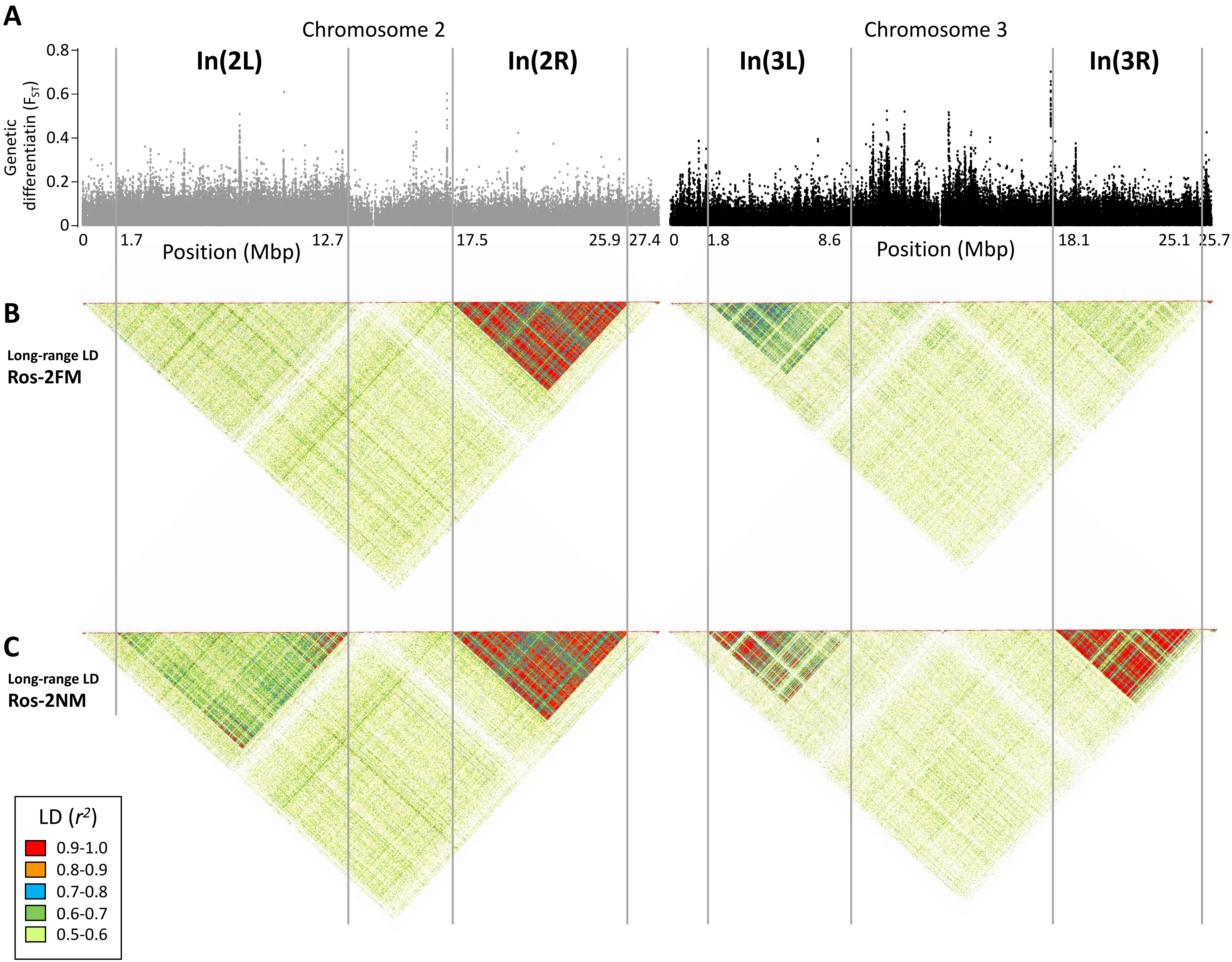


**Supplementary Figure 6 Long range linkage disequilibrium along chromosomes 2 and 3**. The block of slightly elevated genetic differentiation in chromosome arm 2L (**A**) correspond to a block of elevated long-range LD in the NM type (**C**), but not the full moon type (**B**). Elevated long-range LD suggets that the inversion is polymorphic in the repective population, congruent with the genotyping results presented in Figure 2 (the higher the allele frequency of the inverted haplotype, the stronger is LD).
