## Supplementary Figure 7 for "An oligogenic architecture underlying ecological and reproductive divergence in sympatric populations"

**A**

| Tested models for each QTL pair | | |
|---|---|---|
| | Mf = full model | *additive + epistatic effects* |
| | Ma = additive model | *only additive effects* |
| | M1 = one QTL model | *only one of the two QTL exists* |

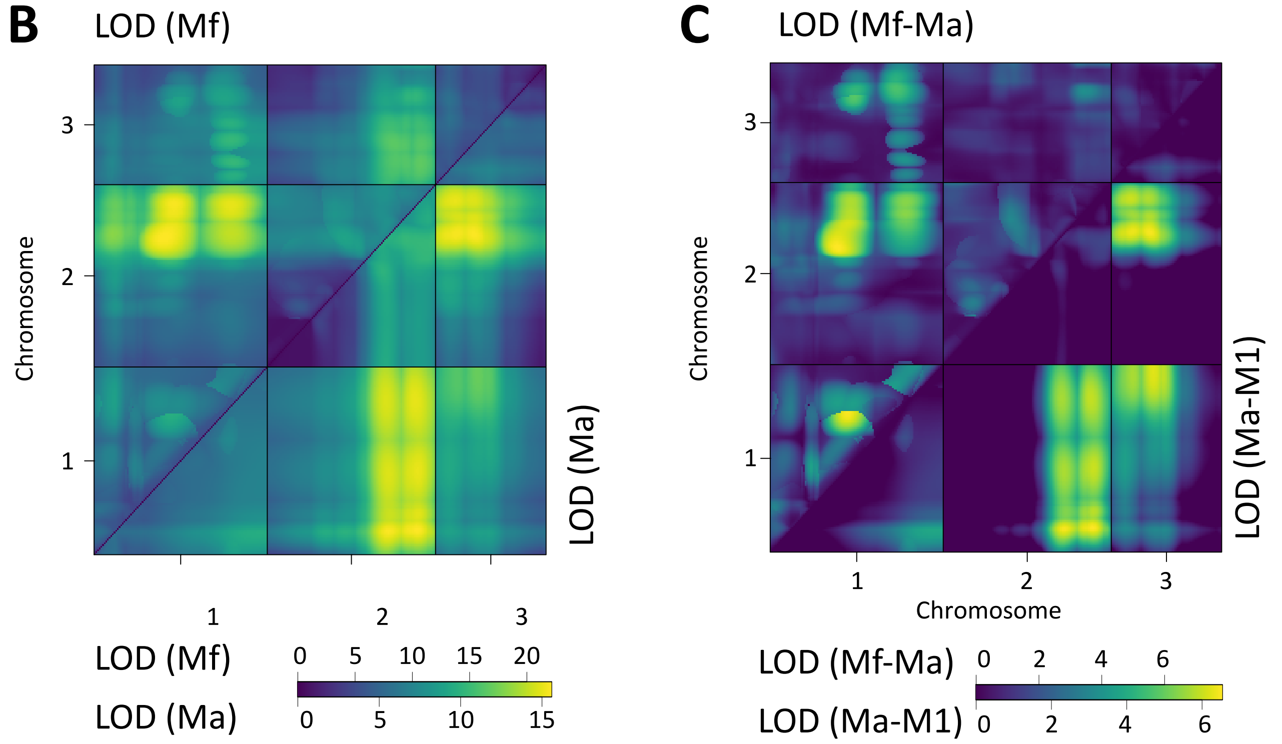


**D**

| Interacting QTL | | | Tests for epistatic interactions | | | | | |
|---|---|---|---|---|---|---|---|---|
| Chr | Pos 1 (cM) | Pos 2 (cM) | LOD(Mf) | p(Mf) | LOD(Mf-M1) | p(Mf-M1) | LOD(Mf-Ma) | p(Mf-Ma) |
| 1:1 | 46 | 78 | **13.9** | **0** | **8.12** | **0.016** | 4.25 | 0.460 |
| 1:2 | 42 | 72 | **22.1** | **0** | **13.09** | **0.000** | **6.56** | **0.018** |
| 1:3 | 84 | 12 | **14.7** | **0** | **9.01** | **0.004** | 2.78 | 0.925 |
| 2:3 | 93 | 12 | **16.6** | **0** | **7.61** | **0.033** | 1.16 | 1.000 |

**E**

| Interacting QTL | | | Tests for additivity | | | |
|---|---|---|---|---|---|---|
| Chr | Pos 1 (cM) | Pos 2 (cM) | LOD(Ma) | p(Ma) | LOD(Ma-M1) | p(Ma-M1) |
| 1:1 | 12 | 100 | **9.62** | **0** | **3.87** | **0.003** |
| 1:2 | 13 | 90 | **15.57** | **0** | **6.53** | **0.000** |
| 1:3 | 101 | 25 | **11.97** | **0** | **6.23** | **0.000** |
| 2:3 | 74 | 23 | **15.49** | **0** | **6.45** | **0.000** |

**Supplementary Figure 7 Scan for interacting QTL**

All chromosmes were scanned for interacting QTLs with the *scantwo* function of R/qtl. (**A**) For each possible QTL combination three models are tested, allowing for either additve and epistatic effects (full model; Mf), only additive effects (Ma) or assuming that one of the two QTL does not exist (M1). (**B**) The LOD score for the full model (Mf) and the additive-only model (Ma) are similar across the genome. (**C**) Congruently, there LOD scores are much lower for the difference between Mf and Ma (upper triangle). The additive model Ma does much better in explaining the data than assuming only one QTL exists (Ma-M1; lower triangle). (**D**,**E**) Statistical tests for the detected QTL combinations. Only in one case the full model is significantly better than the additive-only model (see p(Mf-Ma)), suggesting that epistatic effects are limited to an interaction between the middle of chromosome 1 and the end of chromosome 2 (D). The additive model is significantly better than assuming there is only one QTL (E).
