## Supplementary Figure 8 for "An oligogenic architecture underlying ecological and reproductive divergence in sympatric populations"

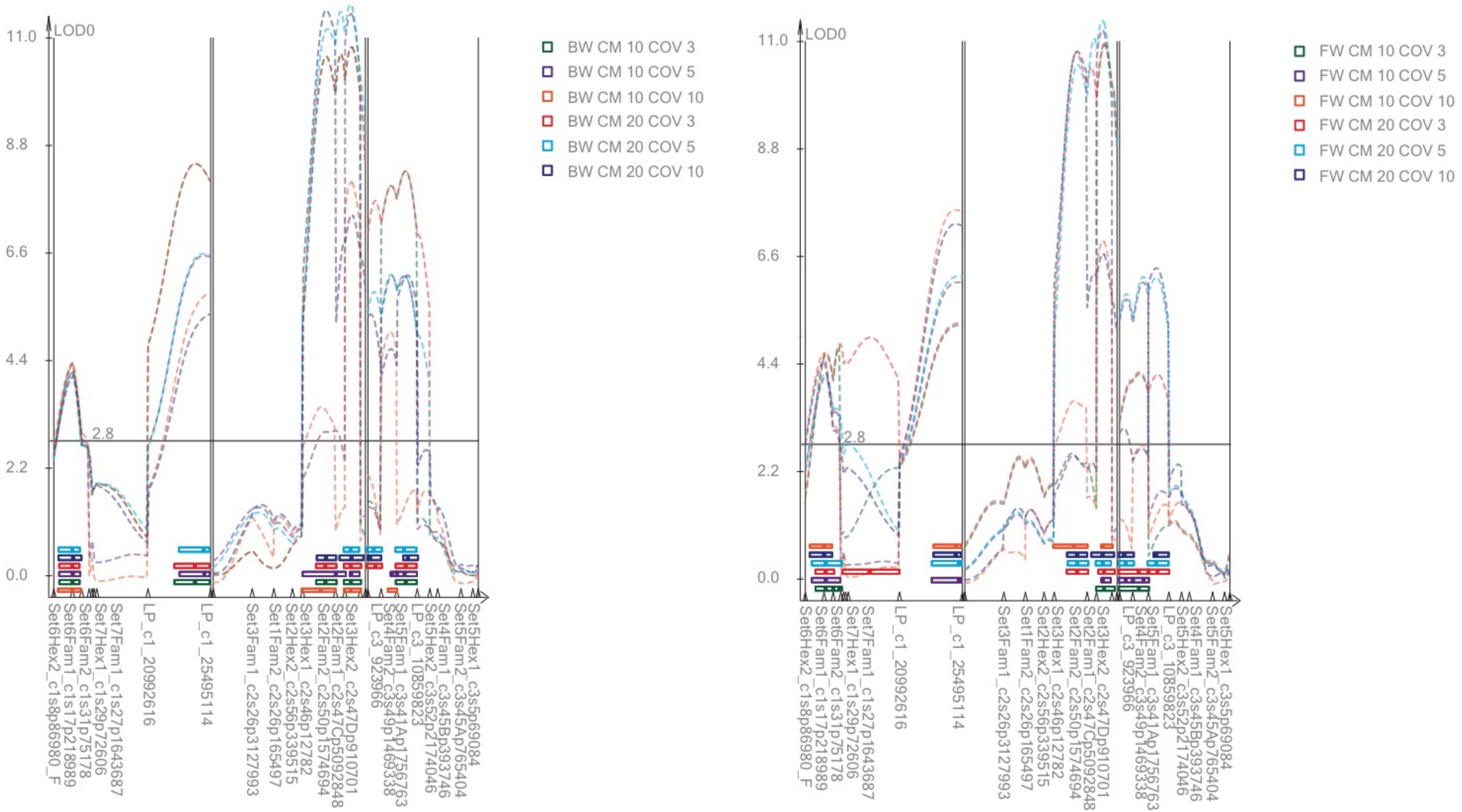


**Supplementary Figure 8 LOD profiles from Composite Interval Mapping (CIM).**
(**A**) Backward selection models. (**B**) Forward selection models. CM = size of the exclusion window in centimorgan; COV = number of covariates. CIM generally identifies the same four QTL as MQM, but sometimes identifies several adjacent QTL in the same genomic region.
