## Supplementary Figure 9 for "An oligogenic architecture underlying ecological and reproductive divergence in sympatric populations"

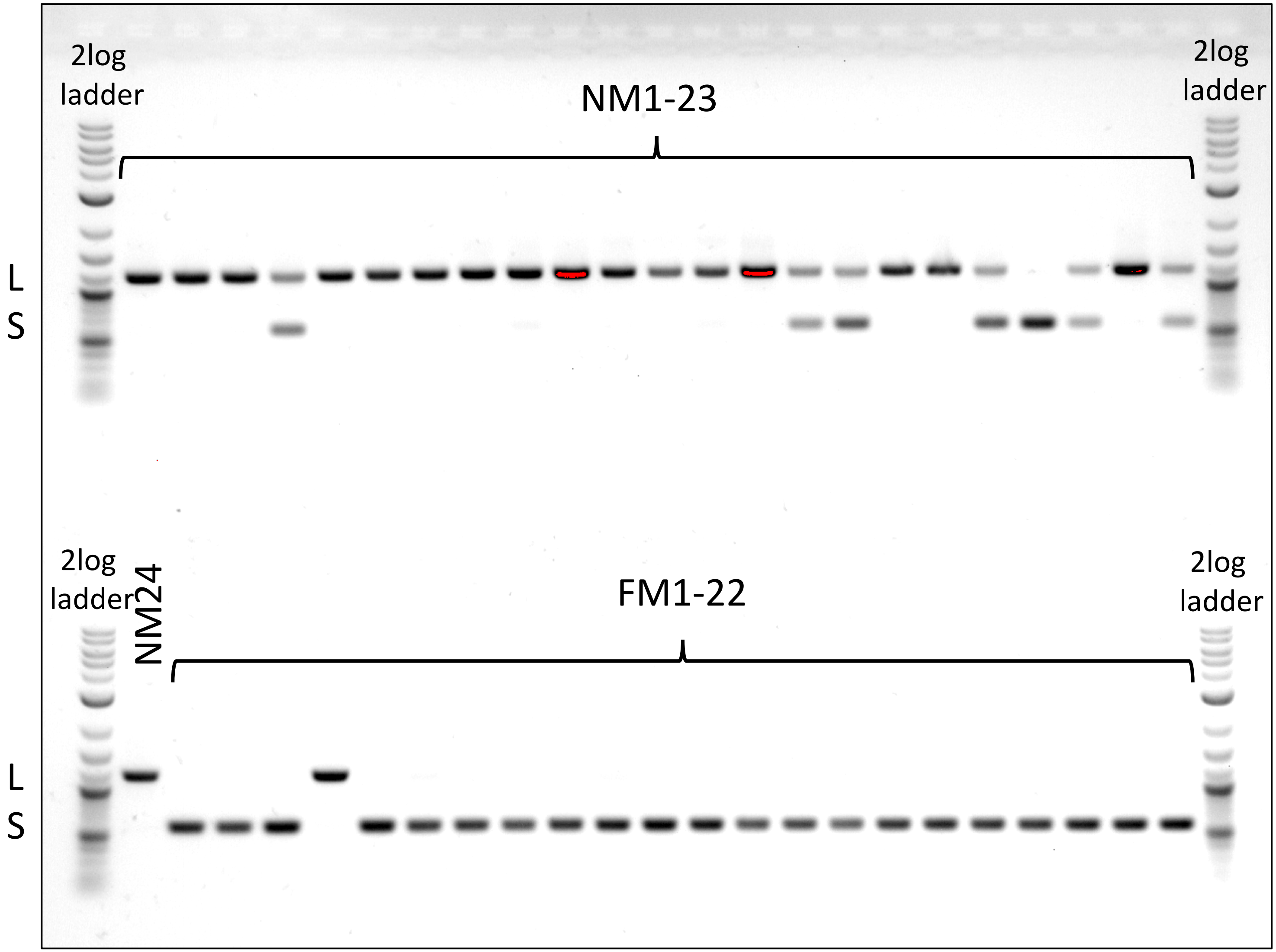


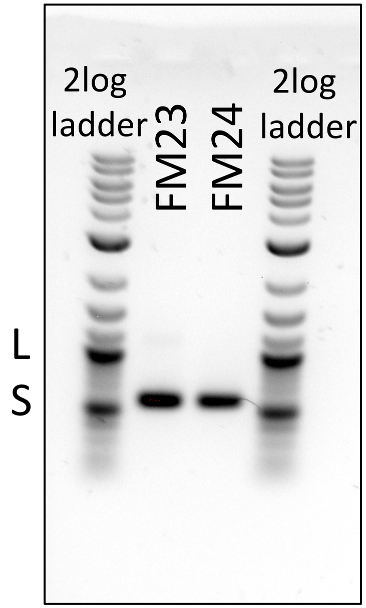


**Supplementary Figure 9 Genotyping of an insertion-deletion in the *period* locus.**
The NM strain is dominated by the long allele (L), the FM strain by the short allele (S).
