## Supplementary Figure 11 for "An oligogenic architecture underlying ecological and reproductive divergence in sympatric populations"

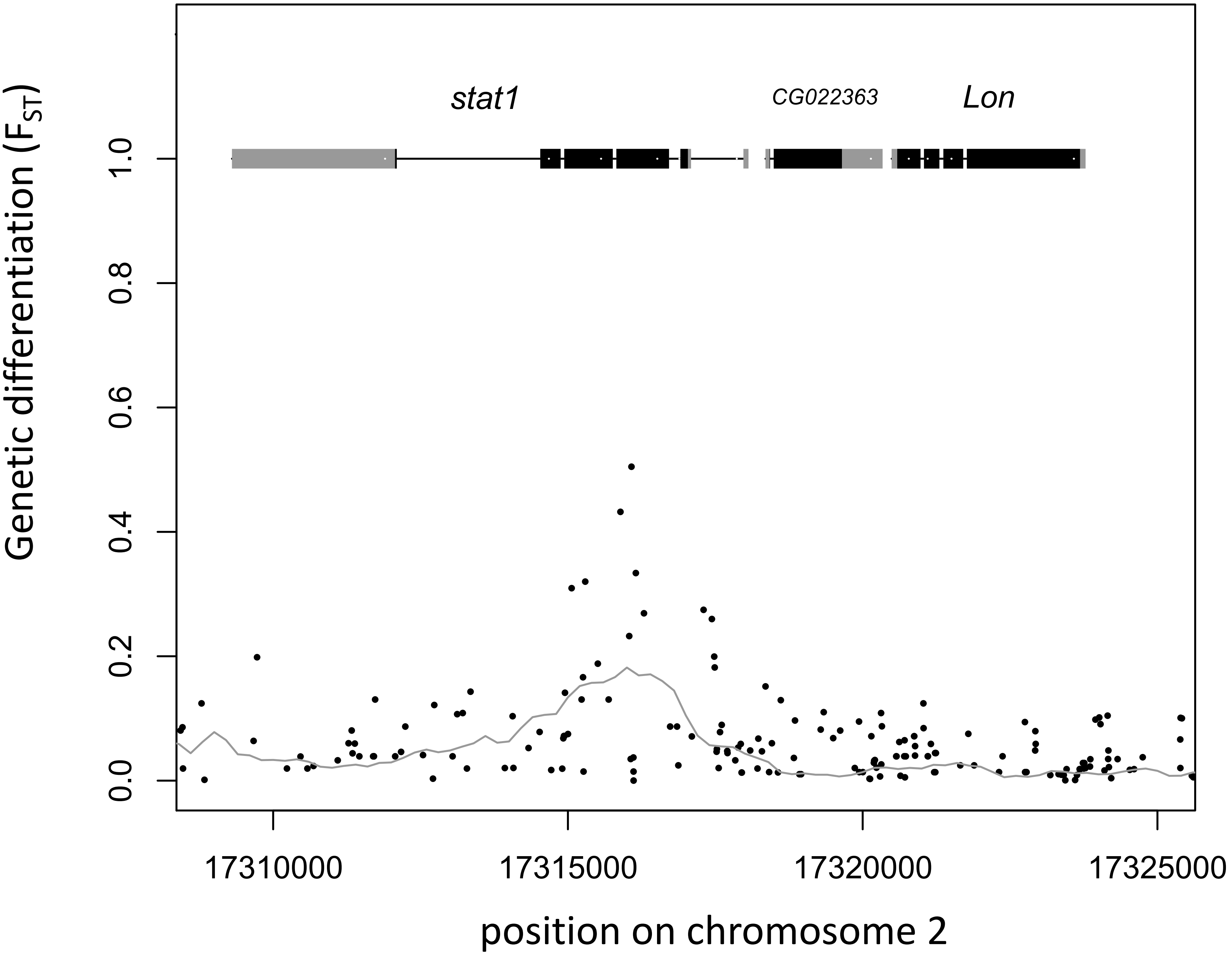


**Supplementary Figure 11 Gene models and genetic differentiation at the *stat1* locus.** Dots are individual SNPs, the line is FST in 1kb sliding windows with 200 bp steps. In the gene models grey boxes are untranslated regions (UTRs), black boxes are coding sequence (CDS).
