## Supplementary Table 1 for "An oligogenic architecture underlying ecological and reproductive divergence in sympatric populations"

**Supplementary Table 1** Inversion breakpoints as estimated from long range LD.

| Inversion | Chr | Left breakpoint (bp) | Right breakpoint (bp) | Size (bp) |
|---|---|---|---|---|
| In(1a) | 1 | 2.741.953 | 18.733.448 | 15.991.495 |
| In(1b) | 1 | 4.353.782 | 11.342.207 | 6.988.425 |
| In(2L) | 2 | 1.675.117 | 12.670.058 | 10.944.941 |
| In(2R) | 2 | 17.530.360 | 25.926.796 | 8.396.436 |
| In(3L) | 3 | 1.806.371 | 8.614.086 | 6.807.715 |
| In(3R) | 3 | 18.144.782 | 25.076.351 | 6.931.569 |
