## Supplementary Table 2 for "An oligogenic architecture underlying ecological and reproductive divergence in sympatric populations"

**Supplementary Table 2** Number of variants in inversion windows

| | Number of variants |
|---|---|
| Chr1 | 261.739 |
| In(1a), left arm | 15.972 |
| In(1b) | 83.456 |
| In(1a), right arm | 92.064 |
| In(1c), right arm | 52.125 |
| Chr2 | 249.948 |
| In(2L) | 114.556 |
| In(2R) | 77.001 |
| Chr3 | 191.892 |
| In(3L) | 53.786 |
| In(3R) | 51.788 |
