## Supplementary Table 3 for "An oligogenic architecture underlying ecological and reproductive divergence in sympatric populations"

| Name | Internal name [1] | Chromosome | Position [2] | Motif | Forward primer | Reverse primer | Size [3] | Dye | Set | Scorable [4] | Marker |
|---|---|---|---|---|---|---|---|---|---|---|---|
| MS101 | MS_c1s8p86980 | 1 | 634807 | AG | TCAAGACGCCAGACAAAAGA | CGCCGCTTCAAATTACTGAC | 415 | HEX | 6 | YES | C1M1 |
| MS102 | MS_c1s15p1802123 | 1 | 1467742 | GAA | GGGTGCGTCACAAAAGTTTC | TGTTTGAGCTTTTCCTGTTGC | 220 | HEX | 8 | NO | - |
| MS103 | MS_c1s29p72606 | 1 | 4419208 | TC | TTGCGATTTTCTGTGGTCTG | TTGGTCTTGACTGGCTGTTG | 213 | HEX | 7 | YES | C1M2 |
| MS104 | MS_c1s59p1429135 | 1 | 5544616 | TG | CGACACATATCACGGAGCAG | TCGGCAAATCTAACCATACGA | 306 | FAM | 7 | YES | C1M3 |
| MS105 | MS_c1s19p129932 | 1 | 6487528 | TC | GCCCTGAAGCATTTCAAAAA | TCCTTGTAGAAAGGCGGAAA | 222 | HEX | 6 | YES | C1M4 |
| MS106 | MS_c1s27p1643687 | 1 | 8266251 | AG | TTTAAGTCAATGCGGAAGAGG | GAACAAAAACCATCGCACAA | 100 | FAM | 7 | YES | C1M5 |
| MS107 | MS_c1s22Bp113007 | 1 | 8553163 | TC | ACTTCGCATGTTTCCCAGAA | CCATGTCATCAACTTCCGTCT | 411 | HEX | 7 | NO | - |
| MS108 | MS_c1s31p75178 | 1 | 14704747 | AG | TGGATTGAGATGGACGTTGA | CTGTTCGAGTTCAAATGGAATG | 305 | FAM | 6 | YES | C1M6 |
| MS109 | MS_c1s17p218989 | 1 | 17462705 | TG | CAAGCATGACACATCAAGCA | CACACAAACCGTATCGGTGA | 124 | FAM | 6 | YES | C1M7 |
| MS110 | MS_c1s42p265009 | 1 | 18183719 | GAT | CACGACGCACCACTTAAGAA | CACAAAATCAGTCGGTGGAA | 329 | FAM | 8 | NO | - |
| LP1 | LP_c1_20992616 | 1 | 20992616 | - | TAGAGCGATCCGCGCTAATA | GATTATCTGTTCCACTTGCGATA | 148/227 | - | - | YES | C1M8 |
| MS111 | MS_c1s44p1006880 | 1 | 22933080 | CT | CCATTTAAGGGATGCGTGAT | CAAGCGAAAGCAATTTCCAT | 420 | HEX | 8 | NO | - |
| LP2 | LP_c1_25495114 | 1 | 25495114 | - | TCAAAATGTGAAGTTTCAAAGCA | TTGGAGCATTTCTTCAATGTTT | 114/156 | - | - | YES | C1M9 |
| MS112 | MS_c1s61p885571 | 1 | 25862224 | GA | CAAAACATCTTTTCGGGTCAC | GGGTTCTAGCAAAGCGAAGA | 177 | FAM | 8 | NO | - |
| **Name** | **Internal name [1]** | **Chromosome** | **Position [2]** | **Motif** | **Forward primer** | **Reverse primer** | **Size [3]** | **Dye** | **Set** | **Scorable [4]** | **Marker** |
| MS113 | MS_c2s20p241395 | 2 | 502108 | AG | TTGGCTATTTGGTTGGTCGT | TTGATGTGTTCATCGCACTG | 244 | HEX | 2 | YES | C2M1 |
| MS114 | MS_c2s13p558035 | 2 | 1242367 | TC | GGGTCGCGTCATGATTCTTA | CCAAACCTGAAAATCCATCC | 219 | HEX | 1 | NO | - |
| MS115 | MS_c2s46p12782 | 2 | 2800433 | TC | TTTGGCGTTGTTCAAGTCTCT | TCGACTCAAGCTCCAAAGTG | 215 | HEX | 3 | YES | C2M2 |
| MS116 | MS_c2s56p339515 | 2 | 5674405 | CAT | TGAATGTCGTGTTTGGGTTAAG | GCAGATCTACAATGAAAGGTAATGG | 360 | HEX | 2 | YES | C2M3 |
| MS117 | MS_c2s26p165497 | 2 | 8425973 | TC | CCTTTTTGCCATCGTTCATT | TGTCGAGGGCATGAATATGA | 149 | FAM | 1 | YES | C2M4 |
| MS118 | MS_c2s26p3127993 | 2 | 11358786 | AG | TGAAACCAAAAGTGGCACAA | TGGAAATCCAAAGTGGAAGAG | 149 | FAM | 3 | YES | C2M5 |
| MS119 | MS_c2s50p1574694 | 2 | 14667091 | TC | TGCTTTGACGCTAGTAAACAGG | AAATGATGAACCGTGACTGTCT | 161 | FAM | 2 | YES | C2M6 |
| MS120 | MS_c2s47Cp5092848 | 2 | 18620830 | CA | TGACCTTTTCAAGGCAAATTAAA | TCTCACGTTTTCGATGATGG | 300 | FAM | 2 | YES | C2M7 |
| MS121 | MS_c2s47Cp2334121 | 2 | 21305073 | AG | GTTTCATCATTTCGCTGTTGC | TCCACAAGCTAAGATGCAATTC | 306 | FAM | 3 | NO | - |
| MS122 | MS_c2s47Ap263174 | 2 | 24127353 | TC | AACGATGAATGGCAATTTGA | ACCCACGCATTTTTGGTAAG | 306 | FAM | 1 | NO | - |
| MS123 | MS_c2s47Dp910701 | 2 | 25271117 | AC | AATTCGCTTTGACTGGTGCT | GCCATTGAGTGGCCAAATAC | 445 | HEX | 3 | YES | C2M8 |
| MS124 | MS_c2s47Fp635401 | 2 | 27069765 | AG | GCAAATTCAAGGGAATCCTG | TTCCAGCAAAGTCTGTTTCG | 461 | HEX | 1 | YES | C2M9 |
| **Name** | **Internal name [1]** | **Chromosome** | **Position [2]** | **Motif** | **Forward primer** | **Reverse primer** | **Size [3]** | **Dye** | **Set** | **Scorable [4]** | **Marker** |
| LP3 | LP_c3_923966 | 3 | 923966 | - | AAACAATGTCAATTGCCAAATA | TGAGACTGATGATAGATTATGTTCCA | 126/195 | - | - | YES | C3M1 |
| MS125 | MS_c3s49p1469338 | 3 | 3109346 | AG | GCGATTAAATGTCGCGATAAG | CGTCAGAAGTCAAAGTGTCAGG | 349 | FAM | 4 | YES | C3M2 |
| MS126 | MS_c3s41Ap1756763 | 3 | 5995719 | TAA | GTGGAAAGCTTTGGCGTAGA | TGCTTCTTGTCTTGCTCAATG | 149 | FAM | 5 | YES | C3M3 |
| LP4 | LP_c3_10859823 | 3 | 10859823 | - | CGCAGCGTTTTTCAATACAA | TTTGATTTYCTGGGGTGTGA | 149/228 | - | - | YES | C3M4 |
| MS127 | MS_c3s48p3645480 | 3 | 12404930 | CT | GCATGCAGATTCGAAAAACA | AGCGCTGGTTGAAAAAGAGA | 270 | HEX | 4 | NO | - |
| MS128 | MS_c3s52p2174046 | 3 | 16511020 | TG | CGACGACTCAGTGGTTATGC | ACTCCCACACCTTTCAATCAA | 422 | HEX | 5 | YES | C3M5 |
| MS129 | MS_c3s45Bp393746 | 3 | 18127054 | AC | CGGGGCGAGTATGTTATCAA | GAATGAAGGCTGGAAAAAGTCA | 151 | FAM | 4 | YES | C3M6 |
| MS130 | MS_c3s45Ap765404 | 3 | 20771652 | TG | GCACGAACGAGTTGTGAATG | GGCAGATCGAAATGTCTTTGT | 301 | FAM | 5 | YES | C3M7 |
| MS131 | MS_c3s5p69084 | 3 | 22737957 | TG | TGCAATCGTACGTTGAACTCTT | TTGCAAACTGACGCAGACTC | 211 | HEX | 5 | YES | C3M8 |
| MS132 | MS_c3s3p743476 | 3 | 25467715 | AC | TGGAATGGTAGCGGGTTAAA | TCGCTCGTTACAAGATGGAA | 436 | HEX | 4 | YES | C3M9 |

[1] Microsatellite (MS) or length polymorphism (LP); chromosme (c), scaffold (s) and position (p) on CLUMA1.0 reference assembly for MS and on CLUMA2.0 assembly for LP

[2] 5' end of forward primer in CLUMA2.0 refernce assembly

[3] expected amplicon size based on CLUM1.0 reference assembly

[4] microsatellite/length polymorphism was PCR-amplifiable and segregated in a scorable pattern in Ros-FMxNM cross
