## Supplementary Table 4 for "An oligogenic architecture underlying ecological and reproductive divergence in sympatric populations"

**Supplementary Table 4** Final QTL model as given by the *fitqtl* function, obtained with multiple imputation, a normal phenotype model and based on 158 observations.

a) Model formula: y ~ Q1 + Q2 + Q3 + Q4

b) Full Model result

| | df | SS | MS | LOD | %var | p(Chi2) | p(F) |
|---|---|---|---|---|---|---|---|
| Model | 8 | 797.6944 | 99.711800 | 26.71958 | 54.10373 | 0 | 0 |
| Error | 149 | 676.6853 | 4.541512 | | | | |
| Total | 157 | 1474.3797 | | | | | |

c) Drop one QTL at a time ANOVA table

| QTL | df | Type III SS | LOD | %var | F value | p(Chi2) | p(F) |
|---|---|---|---|---|---|---|---|
| 1@12.5 | 2 | 93.71 | 4.450 | 6.356 | 10.32 | 0 | 6.36e-05 |
| 1@105.0 | 2 | 153.28 | 7.005 | 10.396 | 16.87 | 0 | 2.48e-07 |
| 2@89.0 | 2 | 301.80 | 12.653 | 20.470 | 33.23 | 0 | 1.17e-12 |
| 3@14.0 | 2 | 159.54 | 7.263 | 10.821 | 17.56 | 0 | 1.42e-07 |

d) Estimated effects

| QTL | Additive effect | | | Dominance effect | | |
|---|---|---|---|---|---|---|
| | est | SE | t | est | SE | t |
| 1@12.5 | 1.00331 | 0.27691 | 3.623 | 0.76487 | 0.39691 | 1.927 |
| 1@105.0 | 1.42628 | 0.24778 | 5.756 | 0.01726 | 0.40408 | 0.043 |
| 2@89.0 | 2.10262 | 0.28000 | 7.509 | 0.06049 | 0.36010 | 0.168 |
| 3@14.0 | 1.53380 | 0.26992 | 5.682 | -0.55051 | 0.35045 | -1.571 |
| Intercept | 5.43907 | 0.18264 | 29.781 | | | |
