## Supplementary Table 5 for "An oligogenic architecture underlying ecological and reproductive divergence in sympatric populations"

### Supplementary Table 5 Candidate genes as obtained by overlap of genes with strongly differentiated variants and QTL regions.

| FST | QTL | Gene ID | Putative gene | Chromosome | Position | SNPs | | | | | | | | | | | | | Indels | | | | | | | | | | | | | | |
|---|---|---|---|---|---|---|---|---|---|---|---|---|---|---|---|---|---|---|---|---|---|---|---|---|---|---|---|---|---|---|---|---|---|
| | | | | | | CDS: non-synonymous | CDS: synonymous | Stop gained | Stop lost | Stop retained | Start lost | Splice site | 3' UTR | 5' UTR | Intron | Non coding transcript | Intergenic | Intragenic | Frameshift | Inframe (conservative) | Inframe (disruptive) | Stop gained | Stop lost | Stop retained | Start lost | Start retained | Splice site | 3' UTR | 5' UTR | Intron | Non coding transcript | Intergenic | Intragenic |
| >0.6 | | CLUMA_CG020013 | NA | chr1 | 4134425 | . | . | . | . | . | . | . | . | . | . | . | 4 | . | . | . | . | . | . | . | . | . | . | . | . | 2 | . | 2 | . |
| >0.6 | | CLUMA_CG019977 | (sp)_putative_Arginine-hydroxylase_NDUFAF5%2C_mitochondrial | chr1 | 4135600 | . | . | . | . | . | . | . | . | . | . | . | 4 | . | . | . | . | . | . | . | . | . | . | . | . | . | . | 2 | . |
| >0.6 | | CLUMA_CG019979 | (nr)_putative_nuclear_pore_complex_protein_DDB_G0274915-like | chr1 | 4142501 | . | . | . | . | . | . | . | . | . | 2 | . | . | . | . | . | . | . | . | . | . | . | . | . | . | . | . | . | . |
| >0.6 | | CLUMA_CG005654 | (pfam)_DUF1676_domain_containing_protein_CLUMA_CG005654 | chr1 | 4457076 | . | . | . | . | . | . | . | . | . | . | . | 1 | . | . | . | . | . | . | . | . | . | . | . | . | . | . | 1 | . |
| >0.6 | | CLUMA_CG005667 | (pfam)_DUF1676_domain_containing_protein_CLUMA_CG005667 | chr1 | 4457076 | . | . | . | . | . | . | . | . | . | . | . | 1 | . | . | . | . | . | . | . | . | . | . | . | . | . | . | 1 | . |
| >0.6 | | CLUMA_CG000851 | (sp)_similar_to_Dynamin-1-like_protein_CLUMA_CG000851 | chr1 | 4714034 | . | . | . | . | . | . | . | . | . | . | . | . | . | . | . | . | . | . | . | . | . | . | . | . | . | . | 1 | . |
| >0.6 | | CLUMA_CG000853 | NA | chr1 | 4714034 | . | . | . | . | . | . | . | . | . | . | . | . | . | . | . | . | . | . | . | . | . | . | . | . | . | . | 1 | . |
| >0.6 | | CLUMA_CG000841 | NA | chr1 | 4817120 | . | 2 | . | . | . | . | . | . | . | . | . | . | . | . | . | . | . | . | . | . | . | . | . | . | . | . | . | . |
| >0.6 | | CLUMA_CG018930 | (sp)_putative_5'-nucleotidase_domain-containing_protein_3 | chr1 | 5674933 | . | . | . | . | . | . | . | . | . | 2 | . | . | . | . | . | . | . | . | . | . | . | . | . | . | . | . | . | . |
| >0.6 | | CLUMA_CG003005 | (mn)_Ionotropic_receptor_60a_(IR60a) | chr1 | 6300137 | . | . | . | . | . | . | . | . | . | . | . | . | . | . | . | . | . | . | . | . | . | . | . | . | . | . | 1 | . |
| >0.6 | | CLUMA_CG003000 | (sp)_similar_to_Limbic_system-associated_membrane_protein_CLUMA_CG003000 | chr1 | 6300137 | . | . | . | . | . | . | . | . | . | . | . | . | . | . | . | . | . | . | . | . | . | . | . | . | . | . | 1 | . |
| >0.6 | | CLUMA_CG005143 | (sp)_putative_Cadherin-87A | chr1 | 6738725 | . | . | . | . | . | . | . | . | . | 1 | . | . | . | . | . | . | . | . | . | . | . | . | . | . | . | . | . | . |
| >0.6 | | CLUMA_CG005226 | (sp)_putative_Hormone_receptor_4 | chr1 | 7036032 | . | . | . | . | . | . | . | . | . | . | . | . | . | . | . | . | . | . | . | . | . | . | . | 1 | 2 | . | . | . |
| >0.6 | | CLUMA_CG005651 | (sp)_putative_Serine_protease_3 | chr1 | 8399023 | . | . | . | . | . | . | . | . | . | . | . | 1 | . | . | . | . | . | . | . | . | . | . | . | . | . | . | 1 | . |
| >0.6 | | CLUMA_CG005612 | (sp)_putative_Toll-like_receptor_7 | chr1 | 8399023 | . | . | . | . | . | . | . | . | . | . | . | 1 | . | . | . | . | . | . | . | . | . | . | . | . | . | . | 1 | . |
| >0.6 | | CLUMA_CG003833 | (mn)_Ionotropic_receptor_7.x8_(IR7.x8)_pseudogene? | chr1 | 8526350 | 5 | . | . | . | . | . | . | . | . | . | . | . | . | . | . | . | . | . | . | . | . | . | . | . | . | . | . | . |
| >0.6 | | CLUMA_CG003841 | (sp)_putative_Ras_GTPase-activating_protein_raskol | chr1 | 8526350 | . | . | . | . | . | . | . | . | . | 25 | . | . | . | . | . | . | . | . | . | . | . | . | . | . | . | . | . | . |
| >0.6 | | CLUMA_CG018849 | NA | chr1 | 9209667 | . | . | . | . | . | . | . | . | . | 1 | . | . | . | . | . | . | . | . | . | . | . | . | 2 | . | 1 | . | . | . |
| >0.6 | | CLUMA_CG002108 | (sp)_related_to_Retinol_dehydrogenase_11 | chr1 | 11354040 | . | . | . | . | . | . | . | . | . | . | . | 2 | . | . | . | . | . | . | . | . | . | . | . | . | . | . | . | . |
| >0.6 | | CLUMA_CG002110 | (sp)_similar_to_Glutathione_S-transferase_1-1_CLUMA_CG002110 | chr1 | 11354040 | . | . | . | . | . | . | . | . | . | . | . | 2 | . | . | . | . | . | . | . | . | . | . | . | . | . | . | . | . |
| >0.6 | | CLUMA_CG024385 | (sp)_putative_Protein_muscleblind | chr1 | 12759391 | . | . | . | . | . | . | . | . | . | . | . | 1 | . | . | . | . | . | . | . | . | . | . | . | . | . | . | . | . |
| >0.6 | | CLUMA_CG001740 | NA | chr1 | 12759391 | . | . | . | . | . | . | . | . | . | . | . | 1 | . | . | . | . | . | . | . | . | . | . | . | . | . | . | . | . |
| >0.6 | | CLUMA_CG002622 | (sp)_putative_Coiled-coil_domain-containing_protein_lobo | chr1 | 13913382 | . | . | . | . | . | . | . | . | . | 1 | . | . | . | . | . | . | . | . | . | . | . | . | . | . | . | . | 1 | . |
| >0.6 | | CLUMA_CG024509 | NA | chr1 | 13913382 | . | . | . | . | . | . | . | . | . | . | . | . | . | . | . | . | . | . | . | . | . | . | . | . | . | . | 1 | . |
| >0.6 | | CLUMA_CG002549 | (sp)_related_to_3-hydroxyisobutyryl-CoA_hydrolase%2C_mitochondrial_CLUMA_CG002549 | chr1 | 14033949 | 2 | 2 | . | . | . | . | . | . | 1 | 2 | . | 1 | . | . | . | . | . | . | . | . | . | . | . | 1 | . | . | . | . |
| >0.6 | | CLUMA_CG022477 | (sp)_related_to_3-hydroxyisobutyryl-CoA_hydrolase%2C_mitochondrial_CLUMA_CG022477 | chr1 | 14035773 | . | . | . | . | . | . | . | . | . | . | . | 1 | . | . | . | . | . | . | . | . | . | . | . | . | . | . | . | . |
| >0.6 | QTL 1 | CLUMA_CG008386 | (mn)_Tachykinin-like_receptor_at_99D_(TkR99D) | chr1 | 15777096 | 3 | . | . | . | . | . | . | . | . | . | . | . | . | . | . | . | . | . | . | . | . | . | . | . | . | . | . | . |
| >0.6 | QTL 1 | CLUMA_CG020610 | (sp)_putative_Syntaxin-1A | chr1 | 16591767 | . | . | . | . | . | . | . | . | . | . | . | . | . | . | . | . | . | . | . | . | . | . | . | . | 5 | . | . | . |
| >0.6 | QTL 1 | CLUMA_CG020824 | (mn)_period_(per) | chr1 | 17280426 | . | . | . | . | . | . | . | . | 7 | 19 | . | 6 | . | . | . | . | . | . | . | . | . | . | . | 2 | 14 | . | 3 | . |
| >0.6 | QTL 1 | CLUMA_CG002364 | (mn)_pickpocket_CLUMA_CG002364 | chr1 | 17281861 | . | 2 | . | . | . | . | 1 | . | . | 1 | . | 9 | . | . | . | . | . | . | . | . | . | . | . | . | . | . | 3 | . |
| >0.6 | QTL 1 | CLUMA_CG002369 | (sp)_similar_to_Dehydrogenase/reductase_SDR_family_member_on_chromosome_X_CLUMA_CG002369 | chr1 | 17285502 | . | 3 | . | . | . | . | . | . | . | . | . | 3 | . | . | . | . | . | . | . | . | . | . | . | . | . | . | . | . |
| >0.6 | QTL 1 | CLUMA_CG002460 | NA | chr1 | 17676854 | . | . | . | . | . | . | . | . | . | . | . | 1 | . | . | . | . | . | . | . | . | . | . | . | . | . | . | . | . |
| >0.6 | QTL 1 | CLUMA_CG002472 | NA | chr1 | 17676854 | 1 | . | . | . | . | . | . | . | . | . | . | 1 | . | . | . | . | . | . | . | . | . | . | . | . | . | . | . | . |
| >0.6 | QTL 1 | CLUMA_CG006144 | (sp)_related_to_Protein_spitz_CLUMA_CG006144 | chr1 | 17770917 | . | . | . | . | . | . | . | . | . | 1 | . | . | . | . | . | . | . | . | . | . | . | . | . | . | . | . | 2 | . |
| >0.6 | QTL 1 | CLUMA_CG023302 | (sp)_related_to_Protein_spitz_CLUMA_CG023302 | chr1 | 17770917 | . | . | . | . | . | . | . | . | . | . | . | . | . | . | . | . | . | . | . | . | . | . | . | . | . | . | 2 | . |
| >0.6 | QTL 1 | CLUMA_CG008012 | (sp)_putative_DNA_topoisomerase_1 | chr1 | 18087445 | . | . | . | . | . | . | . | . | . | . | . | . | . | . | . | . | . | . | . | . | . | . | 4 | . | . | . | . | . |
| >0.6 | QTL 1 | CLUMA_CG008187 | (sp)_similar_to_N-acetylgalactosaminyltransferase_4_CLUMA_CG008187 | chr1 | 18706611 | 1 | . | . | . | . | . | . | . | . | . | . | . | . | . | . | . | . | . | . | . | . | . | . | . | . | . | . | . |
| >0.6 | QTL 2 | CLUMA_CG020204 | (sp)_putative_Nuclear_pore_complex_protein_Nup133 | chr1 | 25510150 | 1 | . | . | . | . | . | . | . | . | . | . | . | . | . | . | . | . | . | . | . | . | . | . | . | . | . | . | . |
| >0.6 | QTL 2 | CLUMA_CG020208 | (mn)_plum \| (sp)_similar_to_Brother_of_CDO | chr1 | 25527322 | . | . | . | . | . | . | . | . | . | 4 | . | . | . | . | . | . | . | . | . | . | . | . | . | . | . | . | . | . |
| >0.4 | | CLUMA_CG017949 | (sp)_similar_to_3-oxoacyl-[acyl-carrier-protein]_reductase_FabG_CLUMA_CG017949 | chr2 | 7452391 | . | . | . | . | . | . | . | . | . | . | . | 2 | . | . | . | . | . | . | . | . | . | . | . | . | . | . | 1 | . |
| >0.4 | | CLUMA_CG017947 | NA | chr2 | 7452391 | . | . | . | . | . | . | . | . | . | . | . | 2 | . | . | . | . | . | . | . | . | . | . | . | . | . | . | 1 | . |
| >0.4 | | CLUMA_CG017952 | (sp)_putative_Sodium/bile_acid_cotransporter_5 | chr2 | 7461980 | . | 2 | . | . | . | . | . | . | . | . | . | . | . | . | . | . | . | . | . | . | . | . | . | . | . | . | . | . |
| >0.4 | | CLUMA_CG017953 | (mn)_Gustatory_receptor_43_(GR43) | chr2 | 7462968 | 1 | 6 | . | . | . | . | . | . | . | 1 | . | . | . | . | . | . | . | . | . | . | . | . | . | . | . | . | . | . |
| >0.4 | | CLUMA_CG009535 | (sp)_related_to_HIV_Tat-specific_factor_1_homolog | chr2 | 7464868 | 1 | 1 | . | . | . | . | . | . | . | 2 | . | 4 | . | . | . | . | . | . | . | . | . | . | . | . | . | . | . | . |
| >0.4 | | CLUMA_CG017954 | NA | chr2 | 7466572 | . | . | . | . | . | . | . | . | . | . | . | 4 | . | . | . | . | . | . | . | . | . | . | . | . | . | . | . | . |
| >0.4 | | CLUMA_CG024356 | NA | chr2 | 7487587 | . | . | . | . | . | . | . | . | . | . | . | 1 | . | . | . | . | . | . | . | . | . | . | . | . | . | . | . | . |
| >0.4 | | CLUMA_CG003411 | NA | chr2 | 7487587 | . | . | . | . | . | . | . | . | . | . | . | 1 | . | . | . | . | . | . | . | . | . | . | . | . | . | . | . | . |
| >0.4 | | CLUMA_CG004602 | (sp)_similar_to_Protein_tramtrack%2C_beta_CLUMA_CG004602 | chr2 | 9578624 | 2 | . | . | . | . | . | . | . | . | . | . | . | . | . | . | . | . | . | . | . | . | . | . | . | . | . | . | . |
| >0.4 | | CLUMA_CG004863 | (sp)_putative_cAMP-specific_3'%2C5'-cyclic_phosphodiesterase | chr2 | 10568233 | . | . | . | . | . | . | . | . | . | . | . | . | . | . | . | . | . | . | . | . | . | . | . | . | 11 | . | . | . |
| >0.4 | QTL | CLUMA_CG015068 | (sp)_putative_Alpha-L-iduronidase | chr2 | 15853287 | 2 | 1 | . | . | . | . | . | . | . | 1 | . | . | . | . | . | . | . | . | . | . | . | . | . | . | . | . | . | . |
| >0.4 | QTL | CLUMA_CG015349 | (sp)_putative_TBC1_domain_family_member_1 | chr2 | 15854557 | . | . | . | . | . | . | . | . | 2 | 1 | . | . | . | . | . | . | . | . | . | . | . | . | . | . | . | . | . | . |
| >0.4 | QTL | CLUMA_CG012971 | (mn)_signal_transducer_and_transcription_activator_Stat92E-like_1_(stat92El1) \| Stat1-like | chr2 | 17316893 | . | . | . | . | . | . | . | . | . | 9 | . | 1 | . | . | . | . | . | . | . | . | . | . | . | . | 3 | . | . | . |
| >0.4 | QTL | CLUMA_CG022363 | NA | chr2 | 17318296 | . | 1 | . | . | . | . | . | . | . | . | . | 1 | . | . | . | . | . | . | . | . | . | . | . | . | . | . | . | . |
| >0.4 | QTL | CLUMA_CG011298 | (sp)_putative_TGF-beta_receptor_type-1 | chr2 | 20700840 | . | . | . | . | . | . | . | . | . | 5 | . | . | . | . | . | . | . | . | . | . | . | . | . | . | . | . | . | . |
| >0.4 | QTL | CLUMA_CG010830 | (sp)_related_to_Galactosylgalactosylxylosylprotein_3-beta-glucuronosyltransferase_P_CLUMA_CG010830 | chr2 | 22370220 | . | . | . | . | . | . | . | . | . | . | . | 1 | . | . | . | . | . | . | . | . | . | . | . | . | . | . | . | . |
| >0.4 | QTL | CLUMA_CG023463 | (sp)_related_to_Galactosylgalactosylxylosylprotein_3-beta-glucuronosyltransferase_P_CLUMA_CG023463 | chr2 | 22370220 | . | . | . | . | . | . | . | . | . | . | . | 1 | . | . | . | . | . | . | . | . | . | . | . | . | . | . | . | . |
| >0.4 | QTL | CLUMA_CG000081 | (sp)_putative_Membralin | chr3 | 1365270 | . | . | . | . | . | . | . | . | . | 2 | . | . | . | . | . | . | . | . | . | . | . | . | . | . | . | . | . | . |
| >0.4 | QTL | CLUMA_CG007484 | NA | chr3 | 7027320 | 2 | 1 | . | . | . | . | . | . | . | . | . | . | . | . | . | . | . | . | . | . | . | . | . | . | . | . | . | . |
| >0.4 | QTL | CLUMA_CG013280 | (sp)_putative_Protein_ECT2 | chr3 | 9308367 | . | . | . | . | . | . | . | . | . | . | . | . | . | . | . | . | . | . | . | . | . | . | . | . | 3 | . | . | . |
| >0.4 | QTL | CLUMA_CG013356 | (sp)_putative_Protein_furry | chr3 | 9647837 | 4 | 6 | . | . | . | . | . | . | . | 2 | . | 1 | . | . | . | . | . | . | . | . | . | . | . | . | 4 | . | . | . |
| >0.4 | QTL | CLUMA_CG013355 | NA | chr3 | 9647837 | . | . | . | . | . | . | . | . | . | . | . | 1 | . | . | . | . | . | . | . | . | . | . | . | . | . | . | . | . |
| >0.4 | QTL | CLUMA_CG013429 | (sp)_putative_Semaphorin-2A | chr3 | 9982618 | . | . | . | . | . | . | . | . | . | . | . | . | . | . | . | . | . | . | . | . | . | . | . | . | 4 | . | . | . |
| >0.4 | QTL | CLUMA_CG013443 | (sp)_putative_ARF_GTPase-activating_protein_GIT2 | chr3 | 10042796 | 1 | . | . | . | . | . | . | . | . | . | . | . | . | . | . | . | . | . | . | . | . | . | . | . | . | . | . | . |
| >0.4 | QTL | CLUMA_CG013513 | (mn)_TRP_channel_transient_receptor_potential_cation_channel_subfamily_M_(Trpm) | chr3 | 10303595 | . | 24 | . | . | . | . | . | . | . | 12 | . | . | . | . | . | . | . | . | . | . | . | . | . | . | . | . | . | . |
| >0.4 | QTL | CLUMA_CG022407 | (sp)_putative_Cytosolic_carboxypeptidase-like_protein_5 | chr3 | 10362891 | . | 1 | . | . | . | . | . | . | . | . | . | . | . | . | . | . | . | . | . | . | . | . | . | . | . | . | . | . |
| >0.4 | QTL | CLUMA_CG013537 | NA | chr3 | 10377569 | . | . | . | . | . | . | . | . | . | . | . | 2 | . | . | . | . | . | . | . | . | . | . | . | . | . | . | . | . |
| >0.4 | QTL | CLUMA_CG013538 | NA | chr3 | 10377569 | . | . | . | . | . | . | . | . | . | . | . | 2 | . | . | . | . | . | . | . | . | . | . | . | . | . | . | . | . |
| >0.4 | QTL | CLUMA_CG013539 | (mn)_Down_syndrome_cell_adhesion_molecule_2_(Dscam2) | chr3 | 10380101 | . | . | . | . | . | . | . | 4 | . | . | . | . | . | . | . | . | . | . | . | . | . | . | . | . | . | . | . | . |
| >0.4 | QTL | CLUMA_CG024000 | (mn)_Aquaporin_big_brain_(bib) | chr3 | 11105680 | . | . | . | . | . | . | . | . | . | 1 | . | . | . | . | . | . | . | . | . | . | . | . | . | . | . | . | 1 | . |
| >0.4 | QTL | CLUMA_CG013720 | (sp)_putative_Neuroligin-like_protein_glit-1 | chr3 | 11105680 | . | . | . | . | . | . | . | . | . | . | . | . | . | . | . | . | . | . | . | . | . | . | . | . | . | . | 1 | . |
| >0.4 | QTL | CLUMA_CG013726 | (sp)_similar_to_Trypsin%2C_alkaline_A_CLUMA_CG013726 | chr3 | 11127429 | 1 | . | . | . | . | . | . | . | . | 1 | . | . | . | . | . | . | . | . | . | . | . | . | . | . | 1 | . | . | . |
| >0.4 | QTL | CLUMA_CG024669 | (nr)_putative_protein_obstructor-E-like | chr3 | 11131133 | . | . | . | . | . | . | . | . | . | . | . | 2 | . | . | . | . | . | . | . | . | . | . | . | . | . | . | . | . |
| >0.4 | QTL | CLUMA_CG018167 | (sp)_similar_to_Trypsin_beta_CLUMA_CG018167 | chr3 | 11131133 | . | . | . | . | . | . | . | . | . | . | . | 2 | . | . | . | . | . | . | . | . | . | . | . | . | . | . | . | . |
| >0.4 | QTL | CLUMA_CG022604 | (sp)_similar_to_Trypsin%2C_alkaline_A_CLUMA_CG022604 | chr3 | 11136276 | 1 | . | . | . | . | . | . | . | . | . | . | . | . | . | . | . | . | . | . | . | . | . | . | . | . | . | . | . |
| >0.4 | QTL | CLUMA_CG013727 | (sp)_similar_to_Lipase_member_H_CLUMA_CG013727 | chr3 | 11138968 | . | . | . | . | . | . | . | . | . | 2 | . | . | . | . | . | . | . | . | . | . | . | . | . | . | . | . | . | . |
| >0.4 | QTL | CLUMA_CG013728 | (pfam)_DUF2236_domain_containing_protein_CLUMA_CG013728 | chr3 | 11141251 | . | . | . | . | . | . | 1 | . | . | 2 | . | . | . | . | . | . | . | . | . | . | . | . | . | . | 3 | . | . | . |
| >0.4 | | CLUMA_CG015788 | (sp)_putative_Zinc_transporter_2 | chr3 | 13221842 | . | . | . | . | . | . | . | 2 | . | 14 | . | 3 | . | . | . | . | . | . | . | . | . | . | . | . | 6 | . | 1 | . |
| >0.4 | | CLUMA_CG015831 | NA | chr3 | 13221842 | . | . | . | . | . | . | . | 6 | . | . | . | . | . | . | . | . | . | . | . | . | . | . | . | . | . | . | . | . |
| >0.4 | | CLUMA_CG015766 | (sp)_similar_to_Chaoptin | chr3 | 13241308 | . | . | . | . | . | . | . | . | 1 | 3 | . | 3 | . | . | . | . | . | . | . | . | . | . | . | . | 1 | . | 1 | . |
| >0.4 | | CLUMA_CG016118 | (nr)_putative_mucin-5AC_CLUMA_CG016118 | chr3 | 13287644 | . | . | . | . | . | . | . | . | . | . | . | 1 | . | . | . | . | . | . | . | . | . | . | . | . | 1 | . | . | . |
| >0.4 | | CLUMA_CG015894 | NA | chr3 | 13291314 | . | . | . | . | . | . | . | . | . | . | . | 1 | . | . | . | . | . | . | . | . | . | . | . | . | . | . | . | . |
| >0.4 | | CLUMA_CG024579 | (sp)_putative_Inositol_monophosphatase_1 | chr3 | 14294877 | . | . | . | . | . | . | . | . | . | 1 | . | . | . | . | . | . | . | . | . | . | . | . | . | . | . | . | . | . |
| >0.4 | | CLUMA_CG015899 | (mn)_ligand-dependent_nuclear_receptor_transcription_factors_ecdysone-induced_protein_78C_(Eip78c) | chr3 | 14313566 | . | . | . | . | . | . | . | . | . | . | . | . | . | . | . | . | . | . | . | . | . | . | . | . | 10 | . | . | . |
| >0.4 | | CLUMA_CG016302 | (nr)_related_to_LOW_QUALITY_PROTEIN:_growth_arrest_and_DNA_damage-inducible_protein_GADD45_alp | chr3 | 15205513 | . | 2 | . | . | . | . | . | 1 | . | . | . | . | . | . | . | . | . | . | . | . | . | . | . | . | . | . | . | . |
| >0.4 | | CLUMA_CG016944 | (sp)_similar_to_Tryptase | chr3 | 15205909 | . | . | . | . | . | . | . | 1 | . | . | . | . | . | . | . | . | . | . | . | . | . | . | . | . | . | . | . | . |
| >0.4 | | CLUMA_CG009336 | NA | chr3 | 18084332 | 1 | . | . | . | . | . | . | . | . | . | . | 13 | . | . | . | . | . | . | . | . | . | . | . | . | . | . | 4 | . |
| >0.4 | | CLUMA_CG009337 | NA | chr3 | 18084693 | 1 | . | . | . | . | . | . | 3 | . | . | . | 13 | . | . | . | . | . | . | . | . | . | . | . | . | . | . | 4 | . |
| >0.4 | | CLUMA_CG022126 | NA | chr3 | 18088307 | . | . | . | . | . | . | . | . | 1 | . | . | . | . | . | . | . | . | . | . | . | . | . | . | . | . | . | . | . |
| >0.4 | | CLUMA_CG009338 | (sp)_putative_ADP-ribosylation_factor_1 | chr3 | 18092454 | . | . | . | . | . | . | . | 1 | . | . | . | . | . | . | . | . | . | . | . | . | . | . | . | . | . | . | . | . |
| >0.4 | | CLUMA_CG022849 | NA | chr3 | 18300253 | 1 | . | . | . | . | . | . | . | . | . | . | . | . | . | . | . | . | . | . | . | . | . | . | . | . | . | . | . |
| >0.4 | | CLUMA_CG009657 | (sp)_related_to_Condensin_complex_subunit_2 | chr3 | 19273849 | . | 1 | . | . | . | . | . | . | . | . | . | . | . | . | . | . | . | . | . | . | . | . | . | . | . | . | . | . |
| >0.4 | | CLUMA_CG006228 | (sp)_putative_Zinc_finger_MIZ_domain-containing_protein_1 | chr3 | 25493088 | . | . | . | . | . | . | . | . | . | . | . | . | . | . | . | . | . | . | . | . | . | . | . | . | 4 | . | . | . |
